# Type I PRMTs and PRMT5 Independently Regulate Both snRNP Arginine Methylation and Post-Transcriptional Splicing

**DOI:** 10.1101/2020.11.18.389288

**Authors:** Maxim I. Maron, Emmanuel S. Burgos, Varun Gupta, Alyssa D. Casill, Brian Kosmyna, Hongshan Chen, Matthew J. Gamble, Charles C. Query, David Shechter

## Abstract

<u>P</u>rotein a<u>r</u>ginine <u>m</u>ethyl<u>t</u>ransferases (PRMTs) methylate histones, splicing factors, and many other nuclear proteins. Type I enzymes (PRMT1-4,6,8) catalyze mono- (Rme1/MMA) and asymmetric (Rme2a/ADMA) dimethylation; Type II enzymes (PRMT5,9) catalyze mono- and symmetric (Rme2s/SDMA) dimethylation. Misregulation of PRMTs in multiple types of cancers is associated with aberrant gene expression and RNA splicing. To understand the specific mechanisms of PRMT activity in splicing regulation, we treated cells with the PRMT5 inhibitor GSK591 and the Type I inhibitor MS023 and probed their transcriptomic consequences. We discovered that Type I PRMTs and PRMT5 inversely regulate core spliceosomal Sm protein Rme2s and intron retention. Loss of Sm Rme2s is associated with the accumulation of polyadenylated RNA containing retained introns and snRNPs on chromatin. Conversely, increased Sm Rme2s correlates with decreased intron retention and chromatin-association of intron-containing polyadenylated RNA. Using the newly developed SKaTER-seq model, comprehensive and quantitative analysis of co-transcriptional splicing revealed that either Type I PRMT or PRMT5 inhibition resulted in slower splicing rates. Surprisingly, altered co-transcriptional splicing kinetics correlated poorly with ultimate changes in alternatively spliced mRNA. Quantitation of retained intron decay following inhibition of nascent transcription revealed that Type I PRMTs and PRMT5 reciprocally regulate post-transcriptional splicing efficiency.

## Introduction

The mammalian genome encodes nine <u>p</u>rotein a<u>r</u>ginine <u>m</u>ethyl<u>t</u>ransferases (PRMTs 1-9; PRMT4 is also known as CARM1). Arginine methylation is critical in regulating signal transduction, gene expression, and splicing (Guccione and Richard 2019; Lorton and Shechter 2019). For instance, a core function of PRMT5, along with its cofactors pICln and MEP50 (also known as WDR77), is regulation of small nuclear ribonucleoprotein (snRNP) assembly (Meister et al. 2001; Boisvert et al. 2002; Meister 2002; Neuenkirchen et al. 2015). This includes both non-enzymatic chaperoning of Sm proteins via PRMT5/pICln following their translation and also post-translational methylation of SmD1, SmD3, and SmB/B’, by PRMT5-MEP50 (Matera and Wang 2014; Paknia et al. 2016). Following their methylation, these Sm proteins are delivered to SMN where they are bound to small nuclear RNAs (snRNAs) in preparation for further processing and eventual nuclear import (Boisvert et al. 2002; Meister 2002; Pellizzoni 2002; Matera and Wang 2014). Disruption of PRMT5 leads to numerous splicing defects, primarily intron retention (<u>r</u>etained <u>i</u>ntrons; RI) and exon skipping (<u>s</u>kipped <u>e</u>xons; SE) (Boisvert et al. 2002; Bezzi et al. 2013; Braun et al. 2017; Fedoriw et al. 2019; Fong et al. 2019; Radzisheuskaya et al. 2019; Tan et al. 2019). While the mechanistic role of Type I PRMTs (PRMT1-4, 6, 8) in splicing is still poorly understood, recent reports demonstrated that there are consequences on SE following Type I PRMT inhibition (Fedoriw et al. 2019; Fong et al. 2019).

RNA splicing can occur during transcriptional elongation or after the transcript has been cleaved and released from chromatin (Bentley 2014; Neugebauer 2019). Although the importance of PRMTs in preserving splicing integrity is clear, whether PRMTs exert their influence over co-or post-transcriptional splicing is still unknown. Previous work has implicated PRMT5 as a regulator of detained introns (DI)—unspliced introns in polyadenylated transcripts (poly(A)-RNA) that remain nuclear and are removed prior to cytoplasmic export (Braun et al. 2017). PRMT5 has also been indirectly shown to regulate post-transcriptional splicing in *Arabidopsis* (Jia et al. 2020). However, both the mechanism of PRMT5’s function and the role of Type I PRMTs in this process remain unclear.

Here, we test whether PRMTs regulate splicing co-or post-transcriptionally and also probe how arginine methylation of snRNP proteins influences this process. As both Type I PRMTs and PRMT5 have been extensively reported to be required for the pathogenicity of lung cancer (Gu et al. 2012; Ibrahim et al. 2014; Avasarala et al. 2015; Sheng and Wang 2016; Li et al. 2019; Zhang et al. 2019)—yet their role in the transcription and splicing of this disease remains largely unstudied—we used A549 human alveolar adenocarcinoma cells (Lieber et al. 1976) as our model. Using GSK591 (also known as EPZ015666 or GSK3203591), a potent and selective inhibitor of PRMT5 (Duncan et al. 2016), and MS023, a potent pan-Type I inhibitor (Eram et al. 2016), we demonstrate that Sm protein symmetric dimethylation and retained introns are inversely correlated with either Type I or PRMT5 inhibition (PRMTi). Loss of Sm protein Rme2s is associated with an accumulation of poly(A)-RNA bound snRNPs on chromatin. Furthermore, assaying co-transcriptional splicing kinetics reveals that either Type I PRMT or PRMT5 inhibition promotes slower splicing and that these changes do not reflect changes in alternatively spliced mRNA. Inhibition of nascent transcription followed by analysis of retained intron decay demonstrates that PRMTs regulate splicing post-transcriptionally and that Type I PRMT inhibition increases post-transcriptional splicing efficiency whereas PRMT5 inhibition decreases efficiency.

## Results

### Type I or PRMT5 inhibition promotes changes in alternative splicing

PRMTs consume S-adenosyl methionine (SAM) and produce S-adenosyl homocysteine (SAH) to catalyze the post-translational methylation of either one or both terminal nitrogen atoms of the guanidino group of arginine (Gary and Clarke 1998). Based on the isoform of their methylated product, PRMTs are classified as Type I, II, or III **(Figure 1a)**. All PRMTs can generate monomethyl arginine (Rme1). Additional methylations are performed using a distributive mechanism where Type I PRMTs catalyze formation of asymmetric N^G^,N^G^-dimethylarginine (Rme2a). Type II PRMTs (PRMT5 and 9) form symmetric N^G^,N’^G^-dimethylarginine (Rme2s). PRMT5 is the primary Type II methyltransferase (Yang et al. 2015). The Type III PRMT7 has only been shown to generate Rme1 (Zurita-Lopez et al. 2012).

**Figure 1.**
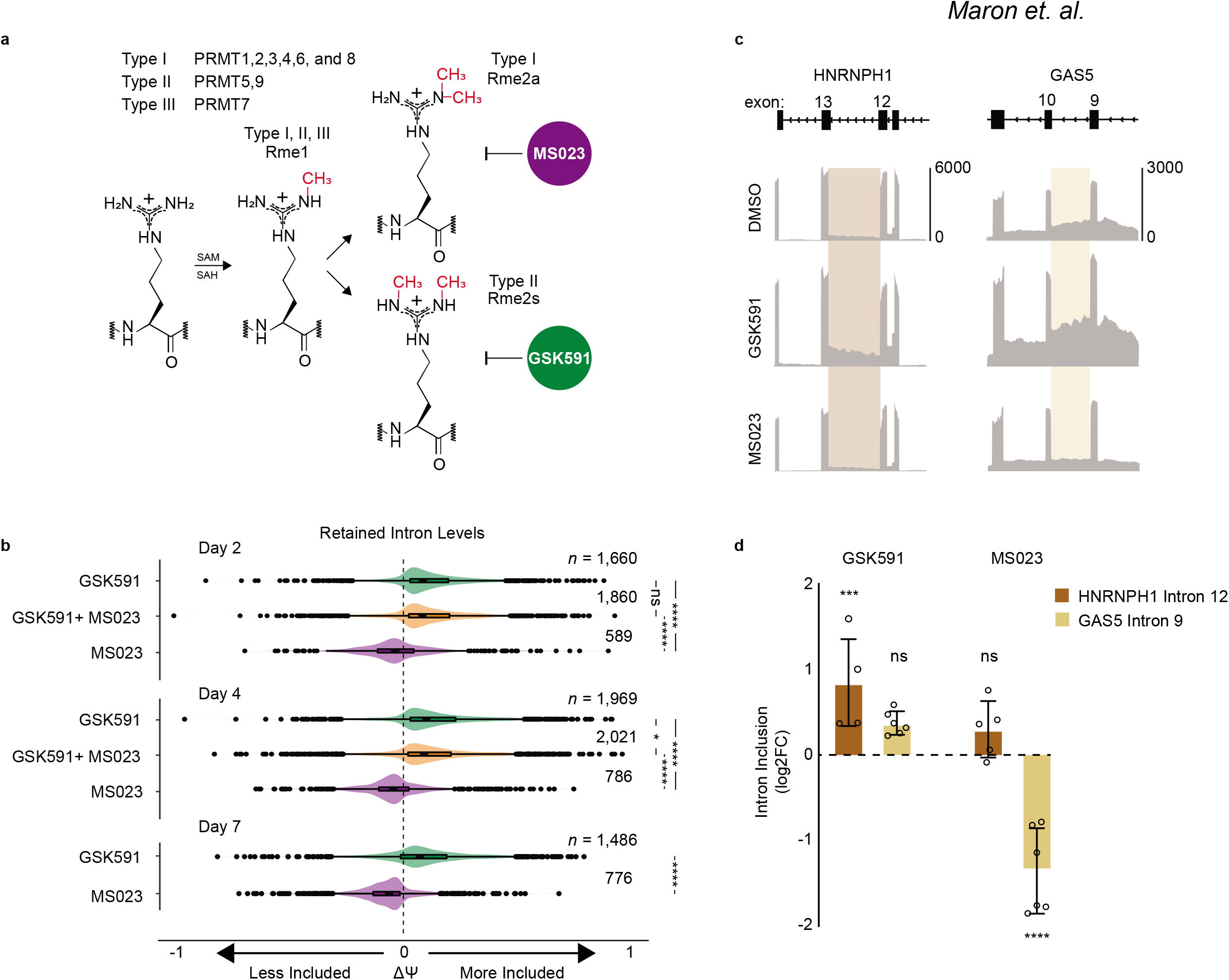
Type I PRMT and PRMT5 activity is required for homeostatic splicing. **a.** Overview of protein arginine methyltransferases and their catalyzed reactions. **b.** Comparison of ΔΨ for RI following PRMTi. ΔΨ = Ψ (PRMTi) – Ψ (DMSO). Treatments are noted as follows: GSK591 (green); MS023 (purple); GSK591 and MS023 (orange). **c.** Genome browser track of poly(A)-RNA seq aligned reads for *HNRNPH1* and *GAS5*. **d.** RT-qPCR validation of RI highlighted in panel f after two-day treatment with DMSO, GSK591, or MS023. Significance determined using two-way ANOVA with Dunnett’s multiple comparisons test (* < 0.05, ** < 0.01, *** < 0.001, **** < 0.0001).

As previous reports indicated that lengthy treatment with PRMTi promotes aberrant RNA splicing, we wanted to determine whether alternative splicing differences with PRMTi occurred as early as day two and, if so, how they evolved over time (Bezzi et al. 2013; Fong et al. 2019; Radzisheus-kaya et al. 2019; Tan et al. 2019). Therefore, we performed poly(A)-RNA sequencing on A549 cells treated with DMSO, GSK591, MS023, or both inhibitors in combination for two-, four-, and seven-days.

Using replicate Multivariate Analysis of Transcript Splicing (rMATS) (Shen et al. 2014) to identify alternative splicing events, at day two we observed significant differences in RI with PRMTi (FDR < 0.05) relative to DMSO **(Figure 1b)**. Whereas RI were increased in GSK591 and cotreatment—signified by a positive difference in <u>p</u>ercent <u>s</u>pliced in (+ΔPSI or Ψ)—RI were decreased in MS023 (-ΔΨ) relative to DMSO. We found that SE were increased in GSK591, MS023, and co-treatment **(Suppl. Figure 1a)**. <u>A</u>lternative <u>5</u>’ <u>s</u>plice <u>s</u>ite (A5SS) and <u>a</u>lternative <u>3</u>’ <u>s</u>plice <u>s</u>ite (A3SS) usage increased only in GSK591 and cotreatment **(Suppl. Figure 1a)**. Meanwhile, <u>m</u>utually e<u>x</u>clusive <u>e</u>xon (MXE) usage was bimodal in all conditions. Generally, the alternative splicing distributions present at day two were propagated through day seven **(Suppl. Figure 1a)**.

To confirm and quantify the presence of PRMT-regulated RI by RT-qPCR, we selected two genes, *HNRNPH1* and *GAS5*. These were identified by rMATS as containing RI at all treatment days **(Figure 1c)**. Using *HNRNPH1* intron 12 and *GAS5* intron 9 primers that flanked the intron-exon boundary and normalized by exon-exon junction primers, at day two following PRMTi we confirmed the presence of the RI **(Figure 1d)**. Consistent with our rMATS data, *HNRNPH1* intron 12 had a significant increase in retention upon GSK591 treatment (*P* < 0.001) and *GAS5* exhibited significantly reduced retention with MS023 (*P* < 0.0001) **(Figure 1d)**.

We next asked whether the RI in our data were common to other datasets in which PRMT activity was perturbed. To accomplish this, we used rMATS on publicly available data (Braun et al. 2017; Fedoriw et al. 2019; Fong et al. 2019; Radzisheuskaya et al. 2019). In these experiments-conducted in a variety of cell lines from diseases including acute myeloid leukemia (THP-1), chronic myeloid leukemia (K562), pancreatic adenocarcinoma (PANC03.27), and glioblastoma (U87)-arginine methylation was inhibited via PRMT knockdown using CRISPRi or with PRMTi including EPZ015666, GSK591/MS023 or GSK591/GSK712. As demonstrated by the high odds ratio (log_2_(OR) > 6) between all the datasets, we showed that there was a highly significant (Fisher’s exact adjusted *P* ≤ 1e-05) overlap in RI **(Suppl. Figure 1b)**. We next intersected all common RI between treatment conditions in A549 across days two, four, and seven **(Figure 2a)**. We found 58 common introns. Strikingly, whereas GSK591 and co-treatment had increased inclusion, for the same introns MS023 resulted in decreased inclusion relative to DMSO **(Figure 2a)**.

**Figure 2.**
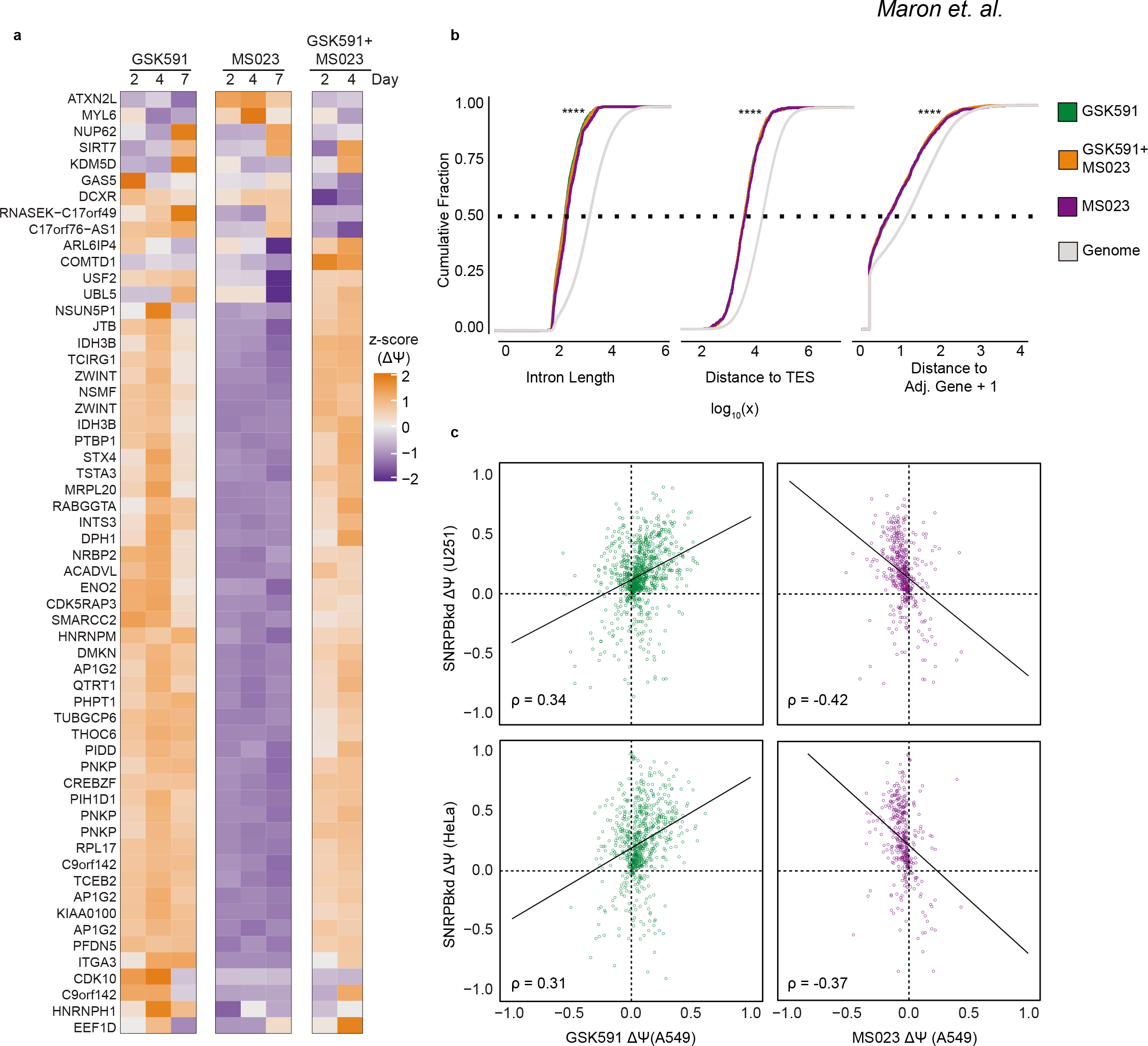
Type I PRMTs and PRMT5 regulate a common group of introns shared with SmB/B’ knockdown. **a.** Heatmap comparing the z-score of ΔΨ for common retained introns in A549 treated with PRMTi across poly(A)-RNA seq experiments on days two, four, or seven. **b.** Cumulative distribution function plots of genomic or RI detected in A549 following two days of PRMTi for intron length, intron distance to the TES, or the distance of the intron containing gene to the nearest adjacent gene in log_10_ units. Significance determined using one-sided Kolmogorov-Smirnov Test (**** < 0.0001). **c.** Comparison of RI ΔΨ detected in day seven of treatment with either PRMTi in A549 or SmB/B’ knockdown in U251 or HeLa cells.

To determine the common characteristics of the RI in A549 treated with PRMTi we analyzed their intra- and inter-gene locations and sequences. We found that the RI were shorter and closer to the transcription end site (TES) compared to the genomic distribution of introns (*P* < 2.2e-16) **(Figure 2b)**. We also observed that RI-containing genes were more likely to be closer to their upstream or downstream genes (*P* < 2.2e-16) **(Figure 2b)**. Moreover, in analyzing the probability of nucleotide distribution at the 5’ and 3’ splice sites we noted both a slight preference for guanine three nucleotides downstream of the 5’ splice site and increased frequency of cytosine in the polypyrimidine tract **(Suppl. Figure 1c)**. This is consistent with previous literature demonstrating that RI have weaker 5’ and 3’ splice sites (Bezzi et al. 2013; Braun et al. 2017; Tan et al. 2019).

As PRMT5 is critical in snRNP assembly, we hypothesized that methylation of the Sm-protein component of snRNPs may play a pivotal role in regulating RI (Friesen et al. 2001a; Friesen et al. 2001b; Meister et al. 2001; Meister 2002; Zhang et al. 2008). We performed rMATS on two publicly available datasets in which SmB/B’ was knocked-down in U251 glioblastoma or HeLa cells (Saltzman et al. 2011; Correa et al. 2016) and compared the RI to those seen with PRMTi. There was a strongly significant (*P* < 2.2e-16) correlation at days two (Spearman’s rank correlation (ρ) = 0.52 or 0.45), four (ρ= 0.44 or 0.39), and seven (ρ = 0.34 or 0.31) with GSK591 treatment in A549 **(Figure 2c)**. Moreover, there was a strongly significant anti-correlation (*P* < 2.2e-16) at days two (ρ = - 0.16 or −0.20), four (ρ= −0.23 or −0.21), and seven (ρ = −0.41 or −0.37) with MS023 treatment **(Figure 2c)**. This result indicated that reduced SmB/B’ protein levels promote an increase in RI that parallels that of PRMTi.

### GSK591 results in Rme2s loss and increased Sm proteins on chromatin

The strong overlap of RI with PRMTi and SmB/B’ knockdown led us to hypothesize that, in A549 cells treated with PRMTi, Sm-protein binding to nascent transcript is compromised. As nascent transcription occurs in the chromatin fraction of cells, we took advantage of the extremely basic isoelectric points (pI) of SmB/B’ (11.2), SmD1 (11.6), and SmD3 (10.33) and performed acid extraction of A549 chromatin treated with DMSO, GSK591, or MS023. Despite performing a stringent wash with 400 mM KCl prior to extraction, GSK591-treated A549 cells had a gross increase of Sm proteins in the acid extracted chromatin fraction **(Figure 3a)**. We did not observe a similar increase in Sm protein levels in the cytoplasmic fractions of either inhibitor treatment **(Suppl. Figure 2a)**. Additionally, by western blot these Sm proteins had reduced Rme2s **(Figure 3a, middle panel)**. Furthermore, consistent with the ability of different classes of PRMTs to scavenge each other’s substrates (Dhar et al. 2013; Lehman et al. 2020), we observed an increase in Rme2a with GSK591. We also noted an increase in Rme2s in the MS023-treated A549 cells **(Figure 3a)**. To further test the large chromatin accumulation of Sm proteins upon GSK591 treatment, we performed an immunofluorescence assay targeting SmB/B’. In GSK591 treated cells, a drastic increase in nuclear DAPI stain overlapping SmB/B’ was apparent **(Figure 3b)**.

**Figure 3.**
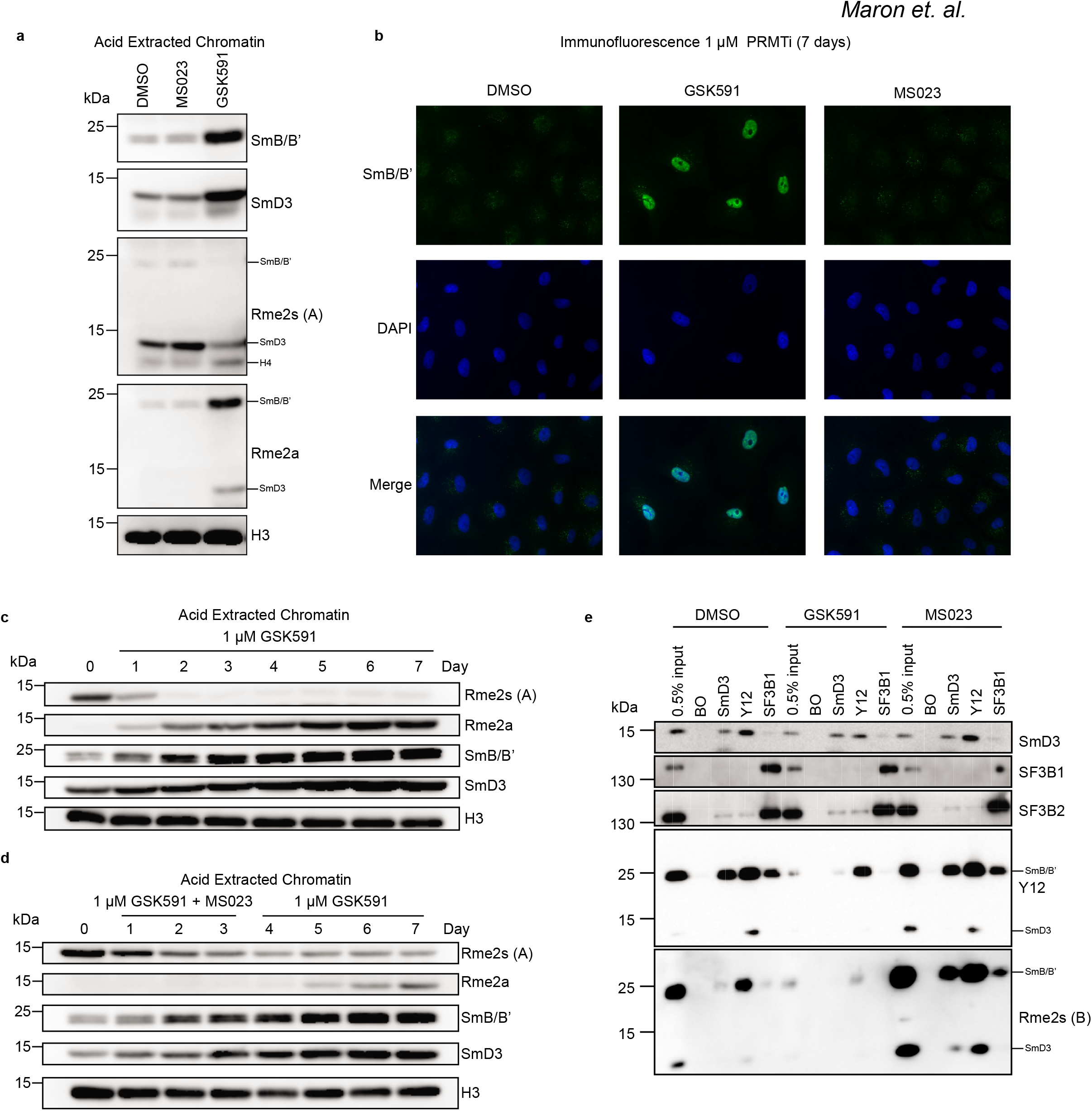
Loss of Rme2s promotes Sm chromatin accumulation. **a.** Western blots of acid extracted chromatin following seven-day treatment with DMSO or PRMTi. **b.** Immunofluorescence for SmB/B’ following seven-day treatment with DMSO or PRMTi. **c-d.** Western blots of acid extracted chromatin with GSK591 treatment for seven days **(c)** or co-treatment PRMTi for three days followed by four days with GSK591 alone **(d)**. **e.** Western blot of co-immunoprecipitation with SmD3, Y12, or SF3B1 antibodies following seven-day treatment with DMSO or PRMTi; BO = beads only.

To assess how Sm protein chromatin association is connected to their arginine methylation, we determined the timeline of Sm Rme2a gain or Rme2 loss in relation to the protein’s chromatin retention. We determined that the majority of Rme2s loss occurs after just one day of PRMT5 inhibition, with almost complete loss after two days **(Figure 3c)**. One day of treatment with GSK591 is associated with a corresponding gain of Rme2a, which increased throughout the seven days of treatment **(Figure 3c)**. The initial accumulation of SmB/B’ and SmD3 was well correlated with the loss of Rme2s and gain of Rme2a. We observed similar results in A549 when using LLY283 (Bonday et al. 2018)—a SAM-analog inhibitor of PRMT5—as well as in IMR90 cells treated with GSK591 **(Suppl. Figure 2b and 2c, respectively)**, signifying that the Rme2s-loss correlated Sm protein chromatin association is specific to PRMT5 inhibition and not unique to A549 cells.

To determine whether accumulation of Sm proteins was due to loss of Rme2s or gain of Rme2a, we performed a co-treatment with GSK591 and MS023. Although the loss of Rme2s follows a similar timeline as GSK591 alone, the gain of Rme2a is delayed until MS023 is removed **(Figure 3d)**. Moreover, the accumulation of Sm proteins on chromatin is coincident with loss of Rme2s and irrespective of Rme2a gain.

### MS023 promotes increased Rme2s on Sm proteins

Although the loss of Rme2s and Sm chromatin accumulation provides insight to a potential mechanism underlying gene expression and splicing changes seen with GSK591, it does not account for the changes seen with MS023. Due to the contrasting effects of Type I PRMT or PRMT5 inhibition on RI ΔΨ, and the common substrate recognition motifs shared by PRMTs, we speculated that an increase in Rme2s due to unopposed PRMT5 activity in MS023-treated cells may underlie the observed changes. We therefore isolated whole cell lysates of A549 cells treated for seven days with either GSK591 or MS023 and-after normalizing for protein concentration-probed the lysates for Rme2s. Whereas Rme2s was completely lost in GSK591 treated cells, in MS023 treated cells there was an increase of Rme2s at molecular weights corresponding to SmB/B’, SmD1, and SmD3 **(Suppl. Figure 2d)**. To confirm that the Rme2s increase was on Sm proteins— typically found in a heptameric ring bound to snRNA, which together which form snRNPs—we performed a co-IP targeting SmD3, Smith Antigen (Y12, recognizing Sm Rme2s) (Lerner and Steitz 1979), and the U2 snRNP factor SF3B1. Indeed, we observed increased Rme2s in the MS023 treated cells **(Figure 3e)**. Additionally, despite the almost complete loss of Rme2s in the input of GSK591-treated cells, even after one week of treatment we observed residual Rme2s on Y12 immunoprecipitated SmB/B’ **(Figure 3e)**. Together these results indicate that Type I or PRMT5 inhibition leads to increased or decreased Rme2s levels of Sm-proteins, respectively.

### Chromatin retained Sm proteins are components of snRNPs

To test if chromatin associated Sm proteins represent intact snRNPs, we first asked whether there was a corresponding increase in snRNA at the chromatin level with GSK591. We treated A549 cells for seven days with DMSO, GSK591, or MS023 and then fractionated them into cytoplasmic, nucleoplasmic, and chromatin components. We then performed northern blotting with [^32^P]-labeled snRNA probes. In GSK591-but not MS023-treated cell nucleoplasm and chromatin fractions, these blots revealed an increased amount of U2, U1, U4, U5, and U6 major snRNAs **(Figure 4a)**. These observations were consistent with intact snRNPs accumulating on chromatin in a PRMT5-inhibited fashion.

**Figure 4.**
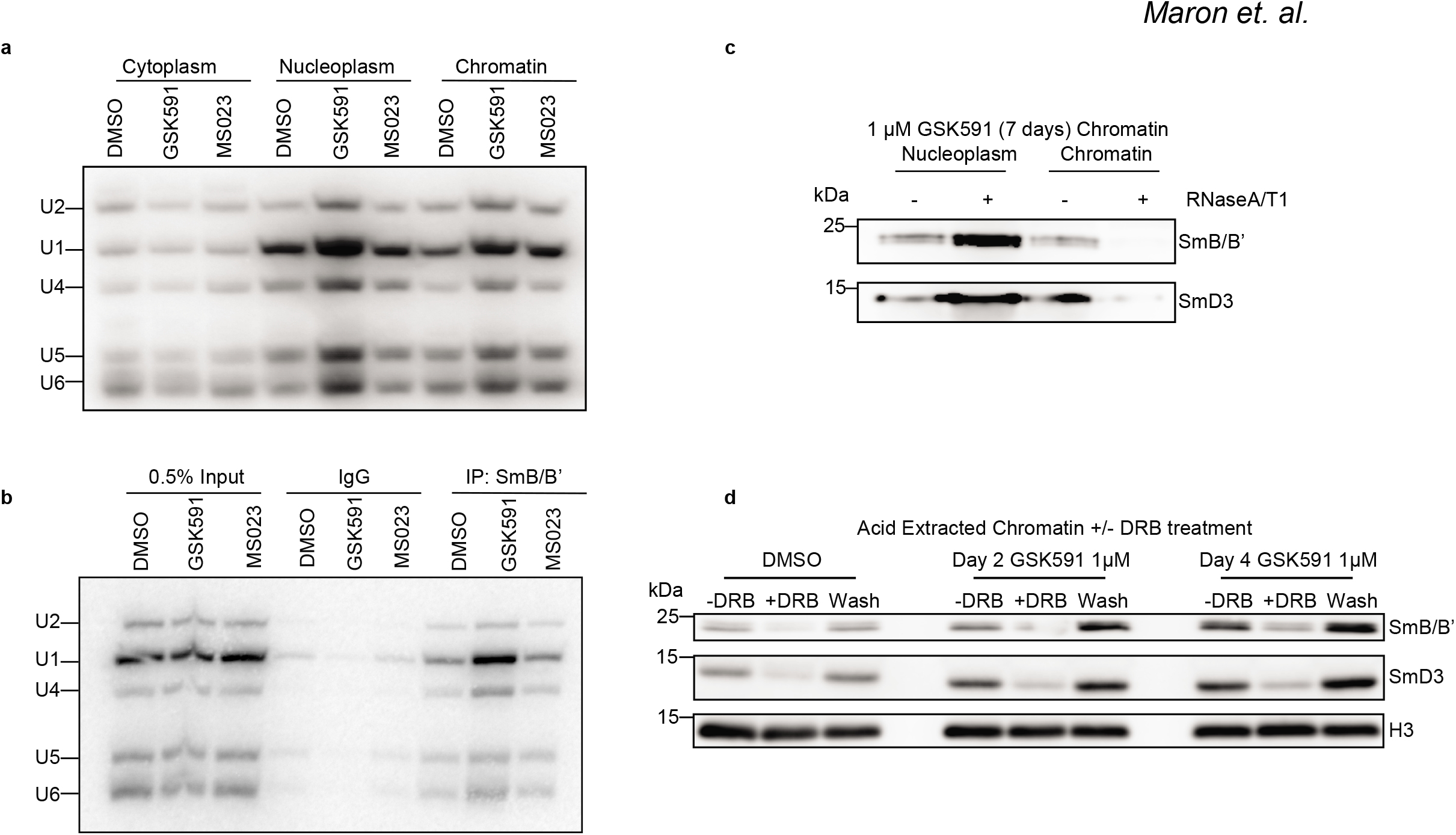
Chromatin-retained Sm proteins are in transcriptionally and RNA-associated snRNPs. **a-b.** Northern blots for snRNA in cytoplasmic, nucleoplasmic, and chromatin fractions following seven-day treatment with DMSO or PRMTi **(a)** or snRNA in RNA immunoprecipitation of SmB/B’ following seven-day treatment of DMSO or PRMTi **(b)**. **c-d.** Western blots of nucleoplasmic and chromatin fractions of control or RNase A/T1 treated chromatin following seven-day treatment with GSK591 **(c)** or acid extracted chromatin in either DMSO or GSK591 treated cells with indicated DRB co-treatment **(d)**.

To determine whether the increased Sm proteins in GSK591-treated cells are in complex with snRNA, using antibodies targeting SmB/B’ we immunoprecipitated snRNPs from whole cell lysate of PRMTi-treated A549 cells and northern blotted with [^32^P]-labeled snRNA probes. We observed that GSK591-treated A549 were associated with increased levels of the five major snRNAs. This was not seen for either DMSO or MS023 treated cells **(Figure 4b)**. We also performed urea polyacrylamide gel electrophoresis followed by SYBR gold staining. We observed increased higher molecular weight RNA species as well as lower molecular weight RNAs corresponding to the snRNAs in GSK591 **(Suppl. Figure 3a)**. This further supported the hypothesis that these Sm proteins represent intact snRNPs. As snRNPs bind to mRNA through complementary base pairing to facilitate splicing, we hypothesized that RNA was the driving force for their chromatin retention. To address whether their chromatin accumulation was RNA-dependent, we isolated the chromatin fraction of A549 cells treated with DMSO, GSK591, or MS023. We then washed the chromatin with 400 mM KCl and, to ensure complete RNA degradation, treated the individual fractions with both RNase A and RNase T1 **(Figure 4c)**. Upon treatment of the nuclear fractions with RNase A/T1, the chromatin retained population of SmB/B’ and SmD3 was shifted to the nucleoplasm, demonstrating that the retained Sm proteins are bound to chromatin through RNA interactions.

### snRNP chromatin accumulation is transcription dependent

As we established that these Sm proteins are intact snRNPs, we asked whether snRNP chromatin accumulation was dependent on active transcription. We treated A549 cells with 100 μM 5, 6-<u>d</u>ichloro-1-β-D-<u>r</u>ibofuranosyl<u>b</u>enzimidazole (DRB)—an inhibitor of p-TEFb that prevents phosphorylation of DSIF and NELF consequently inhibiting RNA pol II pause-release (Wada et al. 1998; Singh and Padgett 2009)—and then probed snRNP chromatin accumulation across time. We noted that 100 μM DRB was able to reduce Sm chromatin association as early as one-hour post-DRB **(Suppl. Figure 3b)**. Next, to determine if this transcriptional elongation-associated Sm chromatin accumulation was dependent upon PRMT5, in GSK591-treated cells, we performed a DRB pulse and washout. Indeed, DRB disrupted snRNP chromatin accumulation **(Figure 4d)**. This indicated that snRNP chromatin accumulation was dependent on transcription of nascent RNA.

### PRMT-dependent changes in co-transcriptional splicing do not reflect changes in steady state mRNA

The dependence of snRNP accumulation on transcription in addition to the modulation of Rme2s with Type I or PRMT5 inhibition prompted us to investigate the kinetics of co-transcriptional splicing. To accomplish this, we used <u>S</u>plicing <u>K</u>inetics and <u>T</u>ranscript <u>E</u>longation <u>R</u>ates by <u>S</u>equencing (SKaTER-seq) (Casill et al. Submitted). As most splicing changes are present as early as two days following GSK591 or MS023 treatment, we used this time point of PRMTi for our analysis. Briefly, this method uses a three-hour DRB treatment to synchronize transcription, followed by a rapid wash-out, to allow productive elongation to commence. Once RNA pol II begins elongating, starting at 10 minutes nascent RNA is collected every five minutes until 35 minutes post-DRB washout **(Figure 5a)**. Nascent RNA is isolated via a 1 M urea wash of chromatin (Wuarin and Schibler 1994) and an additional poly(A)-RNA depletion. The nascent RNA is then sequenced. The rate of nascent RNA formation-including: (1) RNA pol II initiation and pause-release (spawn) rate, (2) elongation rate, (3) splicing rate, and (4) transcript cleavage rate-is then calculated by using a comprehensive model to determine the rates that best fit the sequencing coverage.

**Figure 5.**
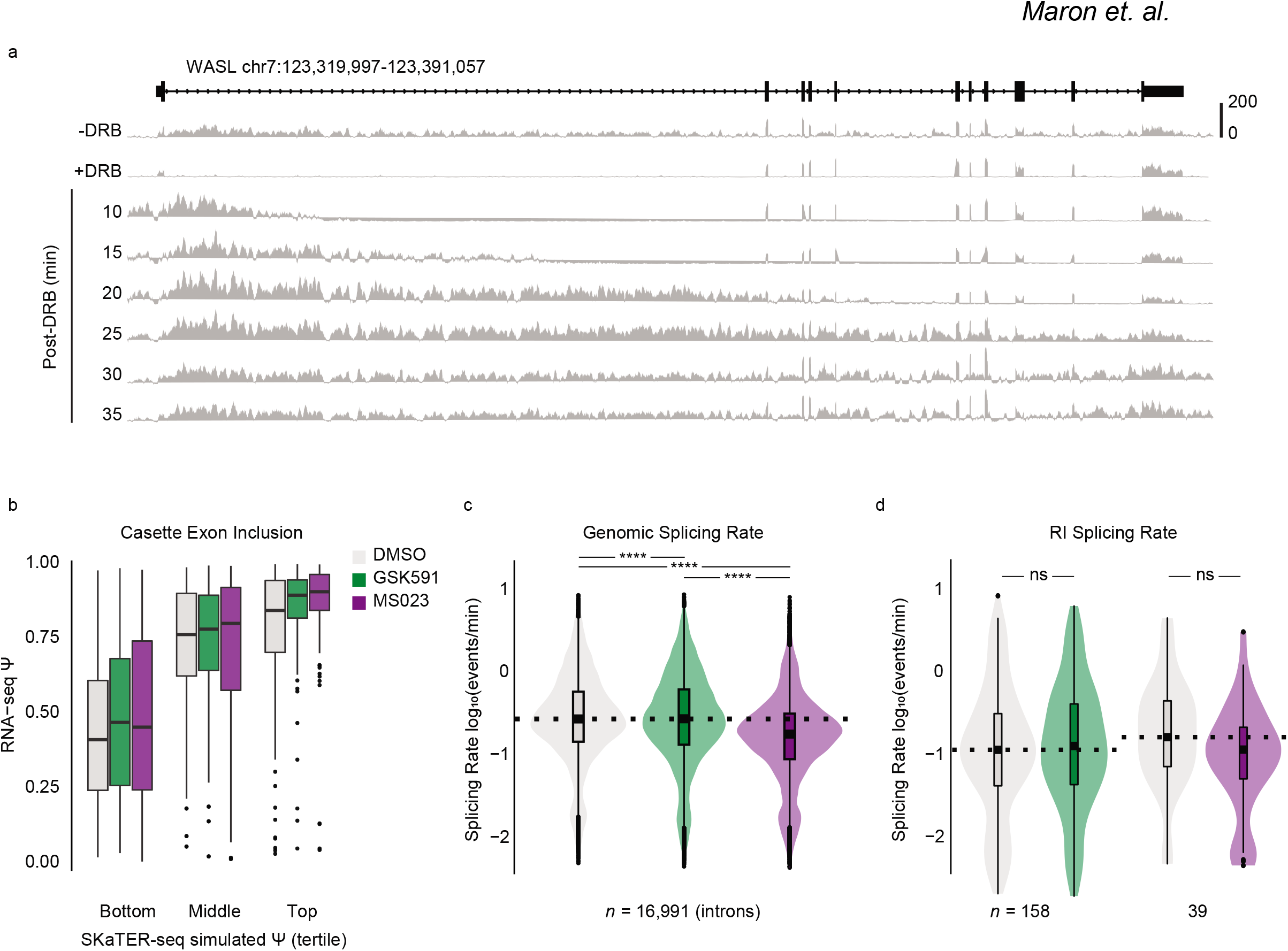
Type I PRMT and PRMT5 activity is necessary for maintaining co-transcriptional splicing kinetics. **a.** Histogram of DMSO-treated SKaTER-seq aligned reads across WASL. **b.** Correlation of SKaTER-seq model simulated cassette exon Ψ versus poly(A)-RNA seq Ψ. **c-d.** Distribution of splicing rates for common genomic introns **(c)** or retained introns **(d)** between DMSO or PRMTi following two days of treatment. Dashed line indicates DMSO median. Solid line within boxplot is condition-specific median. Significance determined using one-sided Kolmogorov-Smirnov Test (ns = not significant, **** < 0.0001).

To assess the accuracy of the rates determined by the SKaTER model, we used the spawn, elongation, splicing, and cleavage rates to simulate a predicted poly(A)-RNA cassette exon Ψ and compared these results with our poly(A)-RNA seq. We successfully predicted cassette exon Ψ detected in poly(A)-RNA in DMSO (ρ = 0.54), GSK591 (ρ = 0.61), and MS023 (ρ = 0.55) (*P* < 2.2e-16) **(Figure 5b)**. Next, we compared spawn rate to poly(A)-RNA seq <u>t</u>ranscripts <u>p</u>er <u>m</u>illion (TPM). As predicted, we observed a strongly significant (*P* < 2.2e-16) correlation between RNA pol II spawn rate and TPM in DMSO, GSK591, and MS023 (ρ = 0.50, 0.51, 0.51, respectively) **(Suppl. Figure 4a)** (Casill et al. Submitted). Consistent with previously published reports, a comparison of splicing rates within each condition confirmed that constitutive introns splice faster than cassette exons **(Suppl. Figure 4b)**. (Pandya-Jones et al. 2013).

As poly(A)-RNA seq revealed opposing alternative splicing changes—specifically increased RI with GSK591 and decreased RI with MS023—we next asked how PRMTi affected the global distribution of splicing rates relative to DMSO. GSK591 resulted in a modest shift toward slower global splicing (*P* = 0.0002, Kolmogorov-Smirnov test statistic (D^-^) = 0.03) and did not significantly change the median of this distribution (adjusted *P* > 0.05 via Dunn’s test) **(Figure 5c)**. Furthermore, the distribution of splicing rates within GSK591 was well correlated with the distribution of the rates in DMSO (ρ = 0.82, *P* < 2.2e-16) **(Suppl. Figure 4c)**. Unexpectedly, MS023 resulted in a significant shift in splicing rates to be slower relative to DMSO (*P* < 2.2e-16, D^-^ = 0.20) **(Figure 5c)**. Despite the drastic decrease in global splicing rates, the distribution of rates within MS023 was strongly correlated to DMSO (ρ = 0.71, *P* < 2.2e-16) **(Suppl. Figure 4c)**. When comparing the splicing rate of RI, we found no significant difference in either GSK591 or MS023 relative to DMSO **(Figure 5d)**.

We next asked how RI splicing rates compared to the genomic distribution of intron rates. We observed that introns that tended to be retained in GSK591 or MS023 were slower to splice (*P* = 0.005, D^-^ = 0.15 and *P* = 0.07, D^-^ = 0.19, respectively) when compared to the genomic distribution in either condition **(Figure 6a)**. When examining additional characteristics of these RI we noted that they had a significantly higher GC percentage in both GSK591 (52%, *P* < 2.2e-16) or MS023 (50%, *P* = 1.87e-5) relative to the global median (41%) **(Figure 6b)**, supporting results from prior reports (Justin et al. 2013; Braunschweig et al. 2014). When comparing transcription rate, RI took less time to transcribe compared to the genomic distribution of introns in both GSK591 and MS023 (*P* = 1.38e-14, D^-^ = 0.42 and *P* = 0.0001, D^-^ = 0.39, respectively) **(Figure 6c)**. As the distribution of RI length was shorter than genomic introns **(Figure 2b)**, we normalized for intron length and determined the intronic elongation rate. Consistent with an increased GC percentage for RI, we found that they have slower elongation rates relative to the genomic distribution **(Suppl. Figure 5a)**. Next, as our total RNA-seq data indicated that RI were more likely to be closer to the TES **(Figure 2b)** we asked whether intron position correlated with splicing rate. Indeed, we determined that introns located closer to the TES tended to have slower splicing rates **(Suppl. Figure 5b)**.

**Figure 6.**
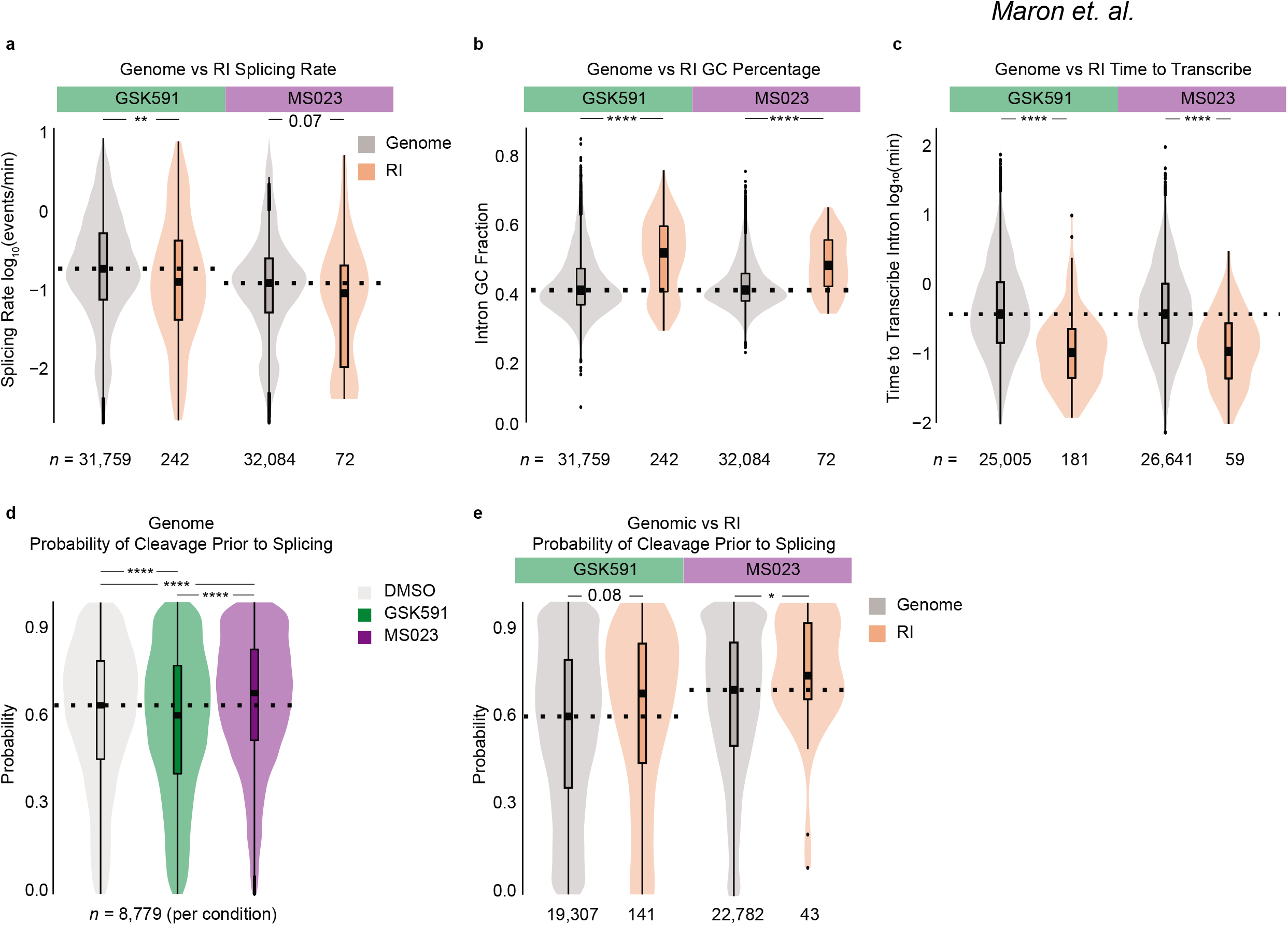
Retained introns have features that decrease the probability of splicing prior to cleavage a-c. Comparison of the distribution of splicing rates **(a)**, GC percentage **(b)**, or time to transcribe intron **(c)** of genomic introns to those of RI in cells treated with PRMTi for two-days. Dashed line indicates global median. Solid line within boxplot is condition-specific median. Significance determined using one-sided Kolmogorov-Smirnov Test (* < 0.05, ** < 0.01, *** < 0.001, **** < 0.0001). **d.** Distribution in probability of cleavage prior to splicing in common introns for cells treated with DMSO, GSK591, or MS023 for two-days. Dashed line indicates DMSO median. Solid line within boxplot is condition-specific median. Significance determined using one-sided Kolmogorov-Smirnov Test (* < 0.05, ** < 0.01, *** < 0.001, **** < 0.0001). **e.** Comparison of the distribution in probability of cleavage prior to splicing for genomic versus RI. Dashed line indicates global median. Solid line within boxplot is condition-specific median. Significance determined using one-sided Kolmogorov-Smirnov Test (* < 0.05, ** < 0.01, *** < 0.001, **** < 0.0001).

### PRMTs regulate splicing post-transcriptionally

The paradox of decreased RI with Type I PRMT inhibition, despite their slower co-transcriptional splicing rates, led us to hypothesize that PRMTs exert their control over splicing post-transcriptionally. To test this hypothesis, we used a metric that considers elongation, splicing, and cleavage rates to determine the probability that a transcript will be cleaved from RNA pol II prior to an intron being spliced. Surprisingly, more than half of all transcripts were likely to be cleaved prior to the completion of splicing **(Figure 6d)**. The global distribution was significantly reduced with GSK591 (*P* < 2.2e-16, D^-^ = 0.07) and increased with MS023 (*P* < 2.2e-16, D^+^ = 0.07) **(Figure 6d)**. Consistent with the hypothesis that PRMTs regulate splicing of RI post-transcriptionally, the probability of transcript cleavage prior to RI splicing was higher when compared to the global distribution in GSK591 (*P* = 0.08, D^+^ = 0.13) or MS023 (*P* = 0.04, D^+^ = 0.28) **(Figure 6e)**. Furthermore, intron position was strongly predictive of whether splicing was likely to occur prior to transcript cleavage: TES proximal introns had a higher probability of cleavage prior to their splicing **(Suppl. Figure 5c)**.

### snRNPs remain bound to poly(A)-RNA

The data from SKaTER-seq demonstrated that for both RI and non-RI the probability of cleavage prior to splicing was greater than 50%. As we discovered that GSK591 or MS023 inversely regulate snRNP arginine methylation and alter their chromatin association dynamics, we asked whether snRNPs are bound to chromatin-associated poly(A)-RNA. To address this question, we used UV-crosslinking (254 nm) followed by cellular fractionation and then poly(A) enrichment of the chromatin fraction. We used a 25-nucleotide competitor poly(A) to ensure the observed interactions were specific. In all conditions, we observed snRNPs bound to poly(A)-RNA **(Figure 7a)**. Furthermore, and consistent with the results described above, in GSK591-treated cells there were more snRNPs on chromatin and bound to poly(A)-RNA. Surprisingly, the poly(A)-associated snRNPs in GSK591 did not contain Rme2s, suggesting that this modification may be dispensable for nuclear import and recognition of nascent RNA. We also noted that the poly(A)-RNAs were associated with H3 and that this interaction was increased in GSK591 and decreased in MS023 **(Figure 7a)**. Importantly, with the addition of the 25-nucleotide competitor poly(A), H3 and snRNP binding to poly(A)-RNA was lost; this confirmed that we had enriched for only poly(A)-associated snRNPs. These observations prompted us to investigate whether candidate RI can be found tethered to chromatin. We detected increased RI in the chromatin fraction of GSK591 treated cells, and depletion of RI in MS023 treated cells **(Figure 7b)**.

**Figure 7.**
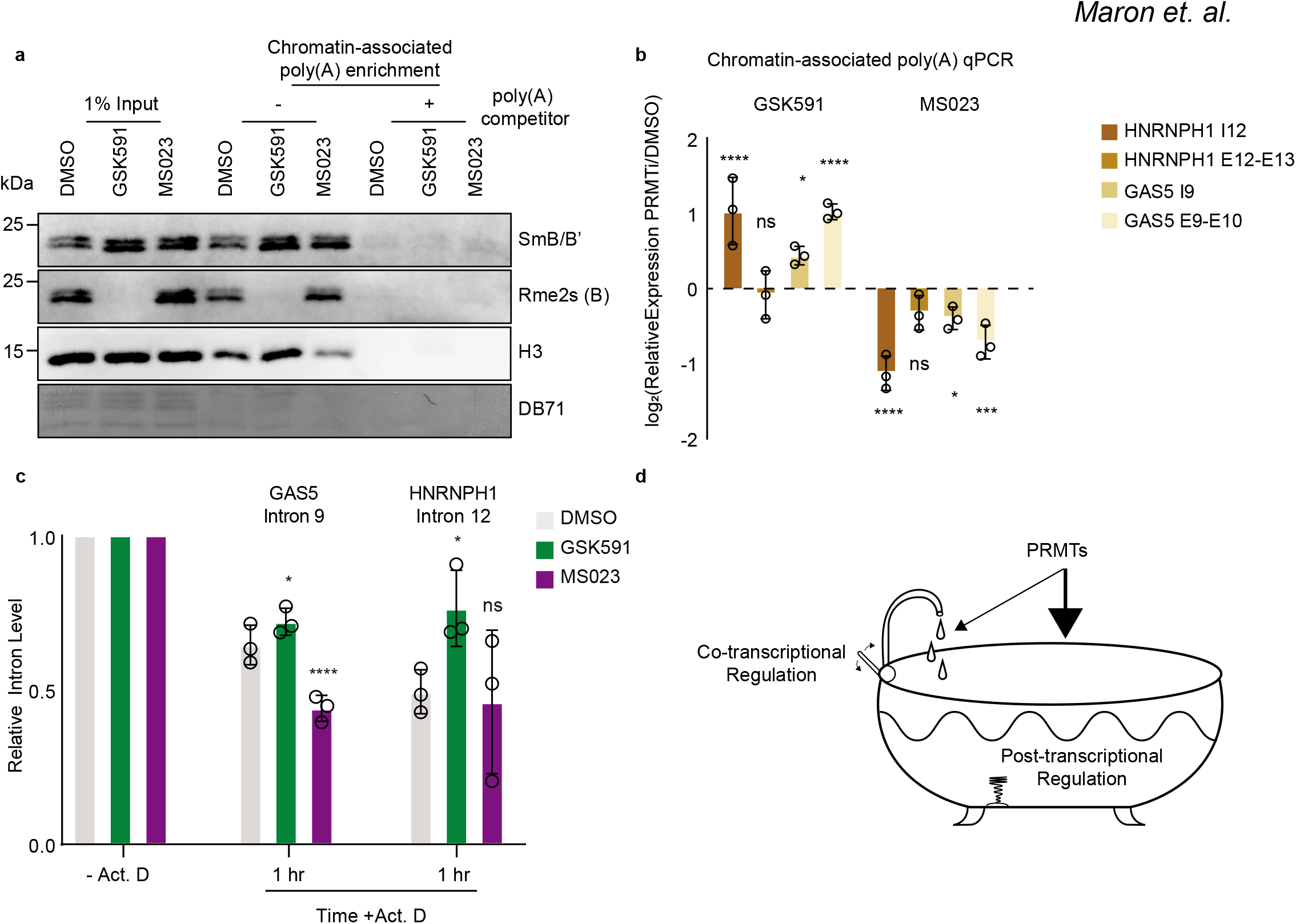
Type I PRMTs and PRMT5 have opposing effects on the removal of retained introns. **a.** Western blot of poly(A)-RNA enriched with oligo(dT) from the chromatin fraction of cells treated with DMSO or PRMTi for two-days with or without the presence of competitor poly(A)-RNA. **b.** RT-qPCR of poly(A)-RNA isolated from the chromatin fraction of cells treated with DMSO or PRMTi for two-days. **c.** Intron decay following treatment with actinomycin D for one hour in cells pre-treated with GSK591 or MS023 for two-days. **d.** Bathtub model representing the regulation of splicing by PRMTs. Nascent transcription is represented by the faucet. The lever (initiation/pause release, elongation, co-transcriptional splicing, cleavage) regulates inflow of water into the bathtub (steady state). Post-transcriptional regulation is represented by the bathtub water (processing/localization) and the drainage (translation/decay).

The slowing of co-transcriptional RNA splicing following PRMTi suggests that there would be an increase in observed RI. Paradoxically, total poly(A)-RNA seq revealed that although RI are increased in GSK591, they are reduced in MS023 relative to DMSO. This led us to hypothesize that MS023 promotes more efficient post-transcriptional splicing, whereas the opposite is true for GSK591. To test this model, we pre-treated A549 cells for two days with GSK591 or MS023 and then blocked transcription using actinomycin D. We then sampled poly(A)-RNA after 60 minutes and probed for *HNRNPH1* intron 12 and *GAS5* intron 9 **(Figure 7c)**. Consistent with a post-transcriptional mechanism, MS023 led to decreased *GAS5* intron 9 when compared to DMSO (*P* < 0.0001), while the removal of *HNRNPH1* intron 12 was not changed. GSK591 led to a significant increase in *GAS5* intron 9 and *HNRNPH1* intron 12 relative to DMSO (*P* < 0.05). We therefore propose the following model: although Type I PRMTs and PRMT5 are required for maintaining the kinetics of co-transcriptional splicing, their impact is ultimately determined post-transcriptionally **(Figure 7d)**.

## Discussion

This study highlights the important role of PRMTs in regulating both snRNP chromatin association and post-transcriptional splicing. We characterized the kinetics of nascent RNA formation and found that, although PRMTi resulted in slower co-transcriptional splicing, these changes were poorly correlated with ultimate alternative splicing changes seen in poly(A)-RNA. We also found that Sm arginine methylation is inversely regulated by Type I PRMTs or PRMT5. Loss of Sm protein Rme2s is associated with a chromatin accumulation of snRNPs as well as transcripts containing RI. When probing post-transcriptional splicing in candidate introns using actinomycin D, whereas GSK591 results in slower intron-decay, MS023 leads to faster decay. The post-transcriptional, rather than co-transcriptional, consequences we observed with PRMTi reflect those seen when we analyzed poly(A)-RNA. Therefore, we conclude that PRMTs regulate post-transcriptional splicing efficiency and transcript-associated snRNP dynamics.

### The effect of PRMT inhibition on splicing

Many reports have documented that PRMT5 has a crucial role in splicing (Bezzi et al. 2013; Koh et al. 2015; Braun et al. 2017; Fedoriw et al. 2019; Fong et al. 2019; Radzisheuskaya et al. 2019; Tan et al. 2019). However, the role of Type I PRMTs in splicing has only recently been appreciated (Fedoriw et al. 2019; Fong et al. 2019). We found that both types of PRMTi carry numerous and diverse consequences on splicing. Remarkably, RI were increased in GSK591, but decreased with MS023 treatment. This is the first report showing that modulating arginine methylation levels can actively enhance splicing. Furthermore, in the presence of co-treatment, alternative splicing changes trended toward that of GSK591 alone, suggesting that Rme2s is dominant in regulating alternative splicing.

Our observation that RI tended to be shorter and closer to the TES as well as enriched in GC content matches data from previous reports (Justin et al. 2013; Braunschweig et al. 2014; Boutz et al. 2015). However, we also observed that genes containing RI were more likely to be closer to their upstream or downstream genes. This suggests a potential for RI to be a consequence of local gene density or higher-order chromatin organization. As has been suggested by others, RI may be an evolutionarily conserved class of introns (Braunschweig et al. 2014; Boutz et al. 2015; Pimentel et al. 2016). This is further supported by our comparison of RI across different publicly available datasets that despite using diverse cell models and methods of inhibiting PRMTs, had a strong overlap with the RI present in our data.

### The effect of PRMT inhibition on snRNP dynamics

Coincident with the changes in splicing, we observed an increase of Sm proteins on chromatin with GSK591. As they co-immunoprecipitate with other Sm proteins and the major snRNAs, we confirmed that these Sm proteins are components of snRNPs. Furthermore, their chromatin association is transcription and RNA dependent. We also showed that the snRNP-chromatin accumulation is coincident with Sm Rme2s and is also consistent with changes in alternative splicing. The overlap of RI following SmB/B’ knockdown in U251 or HeLa cells with either GSK591 or MS023 in A549 suggests that modulation of Rme2s can mimic a loss or gain of function, respectively. Furthermore, a subset of snRNPs are found associated with poly(A)-RNA. Our observation that snRNPs deficient in Rme2s were bound to chromatin-associated poly(A)-RNA suggests that the role of PRMT5 in snRNP ribonucleogenesis is primarily a structural one (Brahms et al. 2001; Meister et al. 2001; Boisvert et al. 2002; Meister 2002). This is supported by the chaperone function of pICln following translation of Sm proteins (Paknia et al. 2016) and successful snRNP assembly *in vitro* by PRMT5-pICln and SMN in the absence of S-adenosyl-L-methionine (Neuenkirchen et al. 2015). Whether the poly(A)-RNA bound snRNPs arise as a result of stalled spliceosomal machinery, intended loading for post-transcriptional splicing, or failed recycling is unclear. Additionally, at which point of nascent transcription these snRNPs are bound and whether they directly influence nascent RNA formation is also unknown.

### The effect of PRMT inhibition on co-transcriptional splicing

By measuring the kinetics of nascent RNA formation using SKaTER-seq, we determined that Type I PRMT or PRMT5 inhibition promotes slower co-transcriptional splicing. Using our measured rates, we were able to successfully predict outcomes of cassette exon usage in poly(A)-RNA. Importantly, predicting cassette exon usage is limited to a few hundred cassette exons and the SKaTER model is designed to measure those spliced co-transcriptionally. Therefore, co-transcriptionally spliced cassette exons will have the most comprehensively measured rates.

GSK591 and MS023 promoted similar co-transcriptional kinetic changes in nascent RNA formation, yet they had contrasting abundance of alternatively spliced isoforms. Consistently, when compared to the global distribution, introns ultimately detected as RI tend to have slower splicing and elongation rates, take less time to transcribe, and are located closer to the TES. This combination is likely to facilitate their retention thus making them amenable to post-transcriptional regulation, as there is less time for splicing to be completed co-transcriptionally. These characteristics were true for Type I PRMT inhibited cells as well, despite having less RI.

Our data reveals that about half of all splicing may not occur prior to transcript cleavage from RNA pol II. Notably, our SKaTER-seq analysis is limited to the number of genes the model was able to solve: roughly 50% of expressed genes that are typically longer genes with fewer isoforms. Nonetheless, our data is consistent with more recent studies highlighting that a considerable fraction of splicing remains to be completed after RNA pol II has transcribed at least a thousand nucleotides past the 3’ SS as well as after transcription has finished (Drexler et al. 2020; Jia et al. 2020; Sousa-Luis et al. 2020).

Although little is known about how post-transcriptional splicing is accomplished, previous work has demonstrated that post-transcriptional spliceosomes are retained in nuclear speckles (Girard et al. 2012). Furthermore, many SR pro-teins—mainly SRSF2—are localized to nuclear speckles (Galganski et al. 2017). Proteomics have indicated that knockdown of Type I PRMTs can increase SRSF2 speckle intensity (Larsen et al. 2016). This presents the possibility that Sm protein Rme2s—which we demonstrated to be increased with Type I PRMT inhibition and decreased with PRMT5 inhibition—and SRSF2 arginine methylation, is required for organization of these interchromatin granules and post-transcriptional splicing. Future work will be needed to characterize the methylarginine dependent mechanisms for successful post-transcriptional splicing.

## Experimental Procedures

### Cell Culture

A549 and IMR90 cells were cultured in DMEM (Corning™, 10-013-CV) and MEM medium (Corning™, 10-009-CV), respectively, supplemented with 10% FBS (Hyclone), 100 μg/mL streptomycin and 100 I.U./mL penicillin (Corning, 30-001-CI) maintained at 37°C with humidity and 5% CO_2_. For this study, fresh cells were purchased from ATCC and tested routinely for *Mycoplasma* (PCR, FWD primer: ACTCCTACGGGAGGCAGCAGT, REV primer: TGCACCATCTGTCACTCTGTTAAC-

CTC).

### Immunofluorescence

Cells were seeded on coverslips (Corning) and allowed to grow in the presence of 0.01% DMSO, 1 μM GSK591 (Cayman), or 1 μM MS023 (Cayman) for seven days. Cells were then washed with 37°C PBS (Hyclone) and fixed with 4% paraformaldehyde at 20-25°C for 10 minutes followed by washing with 4°C PBS. Residual aldehyde was quenched with 0.1 M glycine in 20-25°C PBS for 15 minutes. Permeabilization was performed using 0.1% Triton X-100 at 20-25°C with gentle rotation for 30 minutes. Cover slips were then washed with PBS and blocked for one hour using 0.1% Fish Skin Gelatin in PBS. Primary antibody (Proteintech 16807-1-AP, 1:125) was added to the cover slip and incubated overnight at 4°C in blocking buffer. Coverslips were washed with PBS and incubated with secondary anti-body (Goat antiRabbit Alexa Fluor Plus 488 1:1000) for one hour at 20-25°C followed by PBS wash and mounting with DAPI prolong gold anti-fade (Thermo, P36935). Imaging was performed with an Olympus IX-70 inverted microscope with a 60x objective.

### RT-qPCR

RNA purification was performed using TRIzol (Thermo, 15596026). Isolated total RNA was reverse transcribed with Moloney murine leukemia virus (MMLV) reverse transcriptase (Invitrogen) and oligo-dT primers. LightCycler 480 Sybr Green I (Roche) master mix was used to quantitate cDNA with a LightCycler 480 (Roche). An initial 95°C for 5 minutes was followed by 45 cycles of amplification using the following settings: 95°C for 15 s, 60°C for 1 minute.

### Poly(A)-RNA sequencing

RNA was extracted using RNeasy® Mini Kit (Qiagen, 74104) following the manufacturer’s protocol. RNA quantitation and quality control were accomplished using the Bioanalyzer 2100 (Agilent Technologies). Stranded RNA seq libraries were constructed by Novogene Genetics US. The barcoded libraries were sequenced by Novogene on an Illumina platform using 150nt paired end libraries generating ~30-40 million reads per replicate. Reads were trimmed and aligned to the human genome (hg19) with Spliced Transcripts Alignment to a Reference (STAR) (Dobin et al. 2013). Alternative splicing events were determined using Replicate Multivariate Analysis of Transcript Splicing (rMATS, version 4.0.2) (Shen et al. 2014). IGV (Broad Institute) was used as the genome browser. Graphs pertaining to RNA seq were created using Gviz (Hahne and Ivanek 2016) in R (4.0.2) and assembled in Adobe Illustrator 2020.

### Fractionation of GSK591- and MS023-treated A549 cells via stepwise KCl-elution

After trypsin-digestion, DMSO (0.01%), GSK591- and MS023-treated (1 μM; seven-day exposure) cells were collected. Fractionation was performed as described previously (Shechter et al. 2007). Briefly, cells were resuspended in Hypotonic Buffer (10 mM Tris-HCl pH 8.0 at 4°C, 1.5 mM MgCl_2_, 1 mM KCl, Halt™ Protease and Phosphatase Inhibitor Cocktails (Thermo Scientific™, PI78429 and PI78428, respectively), 1 mM PMSF and 1 mM DTT) and incubated with inversion for 30 minutes at 4 °C. Following centrifugation and removal of Hypotonic Buffer, 0.4-mL <u>N</u>uclear <u>L</u>ysis <u>B</u>uffer (NLB; 10 mM Tris-HCl pH 8.0 at 4°C, 0.1% NP-40, 100-400 mM KCl, Halt™ Protease and Phosphatase Inhibitor Cocktails, 1 mM PMSF and 1 mM DTT) with increasing KCl concentrations was added stepwise to pellets and homogenized for 30 minutes. The remaining pellets were extracted with 0.4 N Sulfuric Acid. The supernatant was isolated and reprecipitated with Trichloroacetic acid solution at a final concentration of 20%. The pellets were then washed 2x with cold acetone, dried under vacuum and resuspended into a cold solution of 0.1% trifluoroacetic acid. Core histones from acid extracted fractions were further analyzed and quantified via HPLC (205 nm).

### Immunoprecipitation

A549 cells were grown with 0.01% DMSO, 1 μM GSK591 (Cayman), or 1 μM MS023 (Cayman) for seven days. Cells were harvested using trypsin (Corning) and washed once with PBS. Cells were then resuspended in NP-40 Lysis Buffer (0.5% NP-40, 50 mM Tris pH 8 at 4 °C, 150 mM NaCl, 1 mM EDTA) supplemented with 40 U/mL RNaseOUT (Thermo, 10777019) and protease inhibitor (Thermo, 78429). Lysates were incubated on ice for 10 minutes followed by sonication 5x for 30 sec on/off on high using a Bioruptor (Diagenode). Lysates were then spun at 10,000 x g for 10 minutes at 4°C. Supernatants were transferred to new low-adhesion RNase-free microcentrifuge tubes and normalized to the same protein concentration. Primary antibody was added followed by incubation overnight at 4°C with gentle rotation. The next morning, Protein A or G agarose (Pierce, 20333 or Millipore-Sigma 16-201, respectively) was equilibrated in lysis buffer and added to the lysates at 4°C with gentle rotation. The beads were washed three times with lysis buffer containing 300 mM NaCl followed by resuspension in either 1x Laemmli buffer for western blotting or TRIzol (Thermo, 15596026) for northern blotting.

### RNase A/T1 treatment and its impact on Sm-protein accumulation onto chromatin

GSK591-treated cells (1 μM for seven-days) were lysed in 1 mL of cold Hypotonic Buffer (as above) at 4°C while continuously inverted onto a tube roller for 30 minutes. After spinning down samples at 10,000 x g for 5 minutes, the pellets were homogenized and resuspended, then split into two equal fractions. To one sample, 0.5 μg of RNase A and 100 Units RNase T1 were added; both samples were kept at 37°C for 20 minutes under continuous mixing using a tube roller.

Supernatants were removed by centrifugation (10,000 g, 5 minutes); matching pellets were subject to a 400-mM KCl wash and extracted via the Sulfuric Acid/TCA method (see above).

### Splicing Kinetics and Transcript Elongation Rates by Sequencing

SKaTER-seq was performed as described (Casill et al. Submitted). Briefly, A549 cells were grown with 0.01% DMSO, 1 μM GSK591 (Cayman), or 1 μM MS023 (Cayman) for two days followed by addition of 100 μM DRB. DRB-containing media was removed, and the cells were incubated at 37°C until the indicated time point. The cells were washed once with 4°C PBS, and lysed by addition of 1 mL CL buffer (25 mM Tris pH 7.9 at 4°C, 150 mM NaCl, 0.1 mM EDTA, 0.1% Triton X-100, 1 mM DTT and protease inhibitor mixture (Thermo 78429)) containing Drosophila melanogaster S2 cell spike-in. Next, lysate was centrifuged at 845 x g for 5 minutes at 4 °C. The pellet resuspended in 1 mL CL buffer without S2 spikein and incubated on ice for 5 minutes. Repeat centrifugation was performed. The supernatant was removed, and cells resuspended in 100 μL GR buffer (20 mM Tris pH 7.9, 75 mM NaCl, 0.5 mM EDTA, 50% glycerol, 0.85 mM DTT) followed by addition of 1.1 mL NL buffer (20 mM HEPES pH 7.6, 300 mM NaCl, 7.5 mM MgCl_2_, 1% NP-40, 1 mM DTT, 1 M Urea). Following a 15-minute incubation, the lysate was spun at 16,000 x g for 10 minutes and the resulting chromatin pellet was resuspended and stored in TRIzol (Thermo, 15596026) at −80 °C. RNA isolation was followed by poly(A)-depletion using the NEBNext Poly(A) mRNA magnetic isolation module (NEB, E7490L). RNA quantitation and quality control were accomplished using the Bioanalyzer 2100 (Agilent Technologies). Stranded RNA-seq libraries were prepared using the KAPA RNA HyperPrep Kit with RiboErase (HMR) and KAPA Unique Dual-Indexed Adapters (Roche) according to instructions provided by the manufacturer. The barcoded paired-end libraries were sequenced by Novogene using a NovaSeq S4, generating ~70 million reads per replicate.

### Chromatin associated poly(A)-RNA enrichment

Poly(A)-RNA isolation was performed with modifications to a previously described protocol (Iadevaia et al. 2018). A549 cells were grown in the presence of 0.01% DMSO, 1 μM GSK591 (Cayman), or 1 μM MS023 (Cayman) for seven days. Cells were washed with 4 °C PBS and irradiated on ice with 100 mJ cm^-2^ in a UV Stratalinker 1800. Cells were centrifuged at 700 x g for 10 minutes at 4 °C. Chromatin was isolated as described above. The chromatin pellet was resuspended in NLB (only with 10 mM Tris pH 7.5 at 4 °C) and sonicated for 5 seconds at 20% amplitude with a probe-tip sonicator using a 1/8” tip. The sonicate was centrifuged at 10,000 x g for 10 minutes and the soluble material transferred to a low-adhesion RNase-free microcentrifuge tube. The samples were split into two separate tubes, one of which received 10 μg of competitor 25-nt poly(A)-RNA. Magnetic oligo-d(T) beads (NEB, S1419S) were equilibrated in NLB and added to the enrichments. The samples were vortexed at 20-25 °C for 10 minutes. The beads were then captured on a magnetic column, and the supernatant transferred to fresh tube for additional rounds of depletion. The beads were washed once with buffer A (10 mM Tris pH 7.5, 600 mM KCl, 1 mM EDTA, 0.1% Triton X-100), followed by buffer B (10 mM Tris pH 7.5, 600 mM KCl, 1 mM EDTA) and lastly buffer C (10 mM Tris pH 7.5, 200 mM KCl, 1 mM EDTA). The RNA was eluted by incubating the beads in 10 μL of 10 mM Tris pH 7.5 at 80 °C for two minutes, capture of magnetic beads using a magnetic column, and quickly transferring the supernatant to a new tube. The beads were then used for two additional rounds of poly(A)-RNA capture.

### Actinomycin D post-transcriptional splicing assay

A549 cells were grown in the presence of 0.01% DMSO, 1 μM GSK591 (Cayman), or 1 μM MS023 (Cayman) for two days. Following a two-day incubation, the media was removed and replaced with media containing 5 μg/μL actinomycin D (Sigma, A1410) with 0.01% DMSO, 1 μM GSK591, or 1 μM MS023 for 60 minutes. RNA was isolated with TRIzol (Thermo, 15596018) and RT-qPCR was performed as described above with ol-igo-d(T) primers.

### Statistical Analysis

All western and northern blots were performed independently at least twice. RT-qPCR was performed at least three-times with independent biological replicates. Statistical analyses were performed using either Prism software (Version 8.3.1, GraphPad) or R (version 4.0.2). To compare distributions, the Kolmogorov-Smirnov (KS) test was used. To account for differences in sample size between global and retained intron (RI) distributions, random sampling from the global population equivalent to the number of RI within the tested condition was performed. This was followed by the KS test and this process was repeated one thousand times after which the median *P* and test statistic was reported. To compare means where greater than two groups exist, either two-way ANOVA with post-hoc Dunnett’s test or the Wilcoxon rank-sum test with post-hoc Dunn’s test was performed. GeneOverlap with Fisher’s exact test was used to determine odds ratios of RI overlap between different RNA seq datasets (Shen 2020).

## Acknowledgments

This work was supported by the National Institutes of Health [R01GM108646 to D.S., GM57829 to C.C.Q., R01GM134379 to M.J.G], the American Lung Association Discovery Award LCD-564723 [D.S] and the HIRSCHL/MONIQUE WEILL-CAULIER TRUSTS to M.J.G.

We thank Dr. Martin Krzywinski (http://mkweb.bcgsc.ca/) for help with data visualization.

## Conflict of Interest

The authors declare that they have no conflicts of interest with the contents of this article. The content is solely the responsibility of the authors and does not necessarily represent the official views of the National Institutes of Health.

## Author contributions

M.I.M. conceived and designed the project, performed biochemical and bioinformatic experiments, interpreted resulting data, and wrote the manuscript. E.S.B. contributed to study design and data interpretation and performed biochemical experiments pertaining to Sm chromatin-association. V.G. performed bioinformatic analysis. A.D.C. performed SKaTER-seq analysis. B.K. and H.C. performed experiments. M.J.G. and C.C.Q. helped with experimental design and data interpretation. D.S. supervised the study and helped with experimental design, data interpretation, and manuscript writing. All authors reviewed the manuscript.

## Supplemental Experimental Procedures

### Antibodies

| Antibody | Vendor | Catalogue | Dilution |
| --- | --- | --- | --- |
| SNRPB | Proteintech | 16807-1-AP | 1:2,000 |
| SNRPD3 | Abcam | ab157118 | 1:2,000 |
| Rme2s (A) | Upstate | Prj-8040 | 1:36,000 |
| Rme2s (B) | CST | 13222S | 1:2,000 |
| Rme2a | Abcam | ab194683 | 1:18,000 |
| H3 | Abcam | ab1791 | 1:100,000 |
| SF3B1 | Bethyl | A300-996A | 1:10,000 |
| SF3B2 | Bethyl | A301-605A | 1:10,000 |
| Y12 | Abcam | ab3138 | 1:2,000 |
| GAPDH | Abcam | ab8245 | 1:10,000 |
| Rabbit IgG HRP | Cytiva | NA934V | 1:100,000 |
| Goat anti-Rabbit 488 | Thermo | 35552 | 1:1,000 |

### Publicly available data used in this study

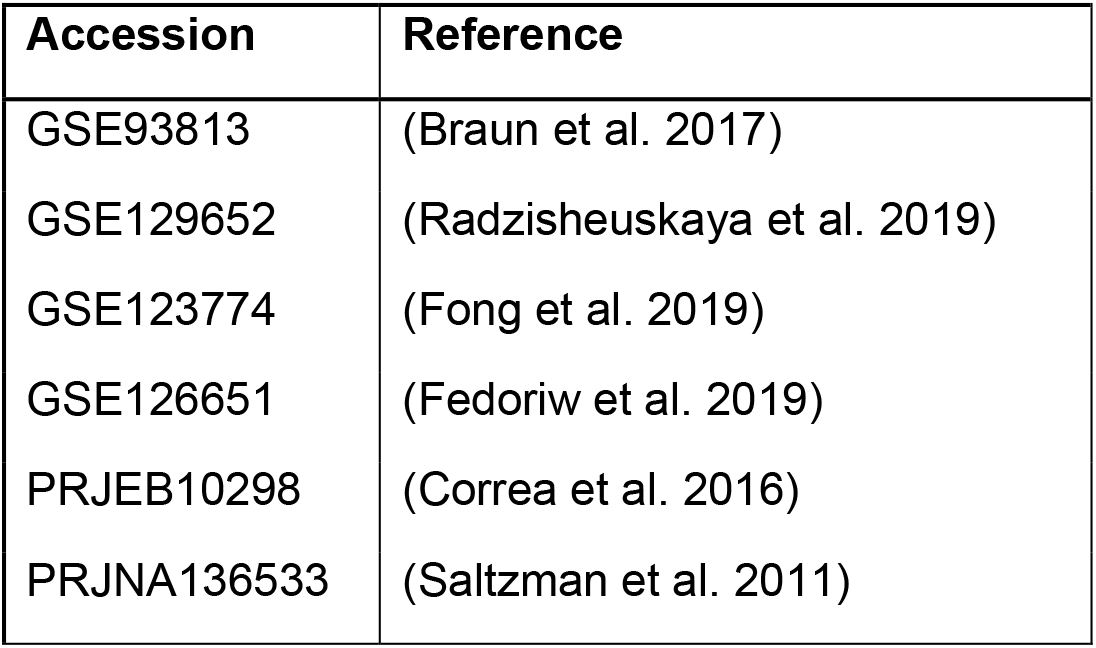

### Primer Sequences

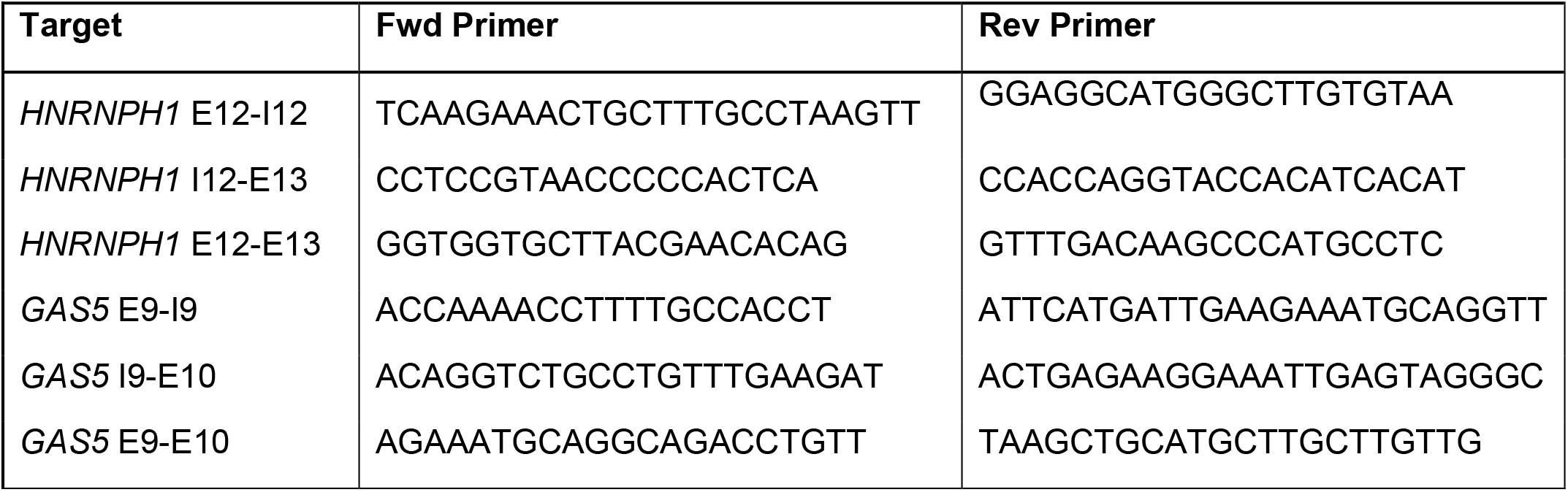

### Western Blotting

For acid extracted chromatin experiments, samples were resolved onto 15% SDS-PAGE (125 V; Tris-Glycine-SDS buffer). Following electrophoresis, gels were equilibrated into cold transfer buffer (15 minutes; Tris-Glycine with 20% Methanol) and further processed for transfer at 4°C (55V, 16h; 0.025% SDS final concentration) onto 0.2 μm Immobilon™-P^SQ^ PVDF membrane (EMD Millipore, ISEQ00010). Following this step, membranes were stained with Direct Blue 71 solution (MeOH/AcOH/water, 50/10/40) and scanned. Membranes were washed with 0.1% PBS-T twice, further blocked with 1% Amersham ECL Prime Blocking Reagent in PBS-T (Cytiva, RPN418) for one hour at 20-25°C. Following a quick wash with 0.1% PBS-T, select membrane segments were incubated with primary antibodies against SmD3, SmB/B’, GAPDH and H3 (overnight; 4°C). Upon processing and incubation with HRP-labelled secondary antibodies (1:50,000; 1h, 20-25°C), membranes were washed, exposed to ECL reagent (Lumigen, TMA-100) and analyzed for their target proteins using an ImageQuant™ LAS 4000 imager. For all other experiments, western blotting was performed as above with the exception that the transfer was performed using 0.45 μm Immobilon™-P PVDF membrane (EMD Millipore, IPVH00010) for one hour at 4 °C at a constant 180 mAmps in 1x NuPAGE buffer (25 mM bicine, 25 mM Bis-Tris, 1 mM EDTA, pH 7.2).

### Northern Blotting

RNA was isolated using TRIzol (Thermo, 15596026) and coprecipitated with GlycoBlue (Thermo, AM9516). The RNA pellet was resuspended in sample buffer (6.8 M Urea in TBE with 10% glycerol and 0.25% Bromophenol Blue/Xylene Cyanide), heated at 90°C for 3 minutes, followed by loading onto an 8% Urea Gel (National Diagnostics) that was pre-run for 45 minutes at 45 watts. The gel was run for one hour at 45 watts in 1x TBE followed by transfer to nitrocellulose in 0.5x TBE at 6V, 130 mAmps for four hours at 4°C. Following transfer, RNA was crosslinked to the membrane at 12 mJ (UV Stratalinker 1800). 5’ end labeling of snRNA probes was performed using ATP [^32^P] (PerkinElmer) with T4 PNK reaction (NEB). Unincorporated ATP [^32^P] was removed using a Microspin G-25 column (Cytiva, 27-5325-01). Post-transfer hybridization was performed in a 37°C water bath overnight with gentle agitation in hybridization buffer containing 100 mM NaHP_4_ pH 7.2, 750 mM NaCl, 1x Denhardt’s Solution (0.02% BSA, 0.02% Ficoll 400, 0.02% Polyvinylpyrrolidone), 1% Herring sperm DNA, and 7% SDS. The next morning, the hybridization solution was carefully discarded according to institutional protocol and the membrane was washed twice with wash buffer (40 mM NaHPO_4_ pH 7.2, 2% SDS, 1 mM EDTA). The wash buffer was also discarded according to institutional guidelines. The membrane was left to expose on a Phosphoimager screen (Cytiva) and imaged at 633 nm using a Typhoon 9400 Variable Mode Imager.

### Fractionation of GSK591- and MS023-treated A549 cells via stepwise KCl-elution

After trypsin-digestion, 1.0 x 10^7^ cells for each DMSO, GSK591- and MS023-treated (1 μM; seven-day exposure) conditions were collected and flash-frozen in liquid nitrogen (long-term storage at −80°C possible). At 4°C, cells were resuspended in 1.5-mL cold Hypotonic Buffer (10 mM Tris-HCl pH 8.0 at 4°C, 1.5 mM MgCl_2_, 1 mM KCl, Halt™ Protease and Phosphatase Inhibitor Cocktails (Thermo Scientific™, PI78429 and PI78428, respectively), 1 mM PMSF and 1 mM DTT); suspensions were continuously inverted using a tube roller for 30 minutes at 4 °C. After spinning down samples at 10,000 x g for 5 minutes, supernatants were isolated and kept at −80°C (Hypotonic Fraction), while remaining pellets were homogenized and washed once with Hypotonic Buffer (30 minutes). Following centrifugation and removal of Hypotonic Buffer, 0.4-mL <u>N</u>uclear <u>L</u>ysis <u>B</u>uffer (NLB; 10 mM Tris-HCl pH 8.0 at 4°C, 0.1% NP-40, 100-400 mM KCl, Halt™ Protease and Phosphatase Inhibitor Cocktails, 1 mM PMSF and 1 mM DTT) with increasing KCl concentrations were added stepwise to pellets (pH kept at 8.0; other additives same as Hypotonic Buffer) and homogenized for 30 minutes. Following the last step (400 mM KCl), pellets were extracted with Sulfuric Acid (0.4 mL, 0.4 N) for 2-3h. Acidic solutions were spun down (20,000 g; 5 minutes), transferred into a new tube and trichloroacetic acid solution (100%) was added for a final concentration of 20%. Samples were vortexed and left on ice for 30 minutes prior to centrifugation. Supernatants were discarded and pellets washed twice with cold acetone. Pellets were dried under vacuum and resuspended into a cold solution of <u>t</u>ri<u>f</u>luoro <u>a</u>cetic acid (TFA, 0.1%), vortexed for one hour and incubated for an extra hour at 4°C. Core histones from acid extracted fractions were further analyzed and quantified via HPLC (205 nm). Following quantification, 7.5 μg of core histones, their matching equivalents in the 100, 200, 300 and 400 mM KCl wash fractions with amounts proportional to the 7.5 μg core histones, and the hypotonic fractions (50 μg total protein) were loaded and resolved onto SDS-PAGE followed by western blotting as described above.

### Splicing Kinetics and Transcript Elongation Rates by Sequencing

A549 (2 x 10^6^ cells/plate) were plated in 100 mm dishes (Corning) containing DMEM supplemented with 10% FBS (Hyclone), 100 μg/mL streptomycin and 100 I.U./mL penicillin (Corning, 30-001-CI) and either 0.01% DMSO, 1 μM GSK591, or 1 μM MS023 such that there was one plate per time point (0, 10, 15, 20, 25, 30, 35 minutes). Cells were incubated at 37 °C with humidity and 5% CO_2_ for two days prior to treatment with DRB. On the day of DRB treatment, all reagents were pre-warmed to 37 °C. Complete media was removed and replaced with complete media containing 100 μM DRB with either 1 μM GSK591 or MS023 (or 0.01% DMSO for negative control). The plates were then incubated at 37 °C with humidity and 5% CO_2_ for three hours. Following DRB incubation, complete media containing 1 μM of either GSK591 or MS023 was prepared at 37 °C. DRB-containing media was removed (except for 0 minute DRB treatment and negative control) and cells were quickly washed twice with 37 °C PBS prior to addition of 6 mL 37 °C complete media containing 1 μM GSK591 or MS023. Cells were then incubated at 37°C until the indicated time point, at which point the plate was removed from the incubator and placed immediately on ice. The supernatant was quickly removed and the cells were washed once with 4°C PBS, followed by addition of 1 mL CL buffer (25 mM Tris pH 7.9 at 4°C, 150 mM NaCl, 0.1 mM EDTA, 0.1% Triton X-100) containing *Drosophila melanogaster* S2 cell spike-in supplemented with 1 mM DTT and protease inhibitor complex (Thermo, 78429). Cells were isolated using a cell scraper and kept on ice until all time points were collected. Following collection of all additional time points, cells were centrifuged at 845 x g for 5 minutes at 4 °C. The supernatant was removed, and the cells resuspended again in 1 mL CL buffer without S2 spikein and incubated on ice for 5 minutes. Cells were then spun again at 845 x g for 5 minutes at 4 °C. The supernatant was removed, and cells resuspended in 100 μL GR buffer (20 mM Tris pH 7.9, 75 mM NaCl, 0.5 mM EDTA, 50% glycerol) supplemented with 0.85 mM DTT followed by addition of 1.1 mL NL buffer (20 mM HEPES pH 7.6, 300 mM NaCl, 7.5 mM MgCl_2_, 1% NP-40) supplemented with 1 mM DTT and 1 M Urea. The lysate was incubated on ice for 15 minutes with vortexing every three minutes. The lysate was spun at 16,000 x g for 10 minutes and the resulting chromatin pellet was resuspended in TRIzol (Thermo, 15596026) (long-term storage at −80°C possible). RNA was isolated per recommended TRIzol protocol followed by poly(A)-depletion using the NEB-Next Poly(A) mRNA magnetic isolation module (NEB, E7490L). Briefly, TRIzol extracted RNA was resuspended in 50 μL RNase-free H2O (Thermo, 10977015). Oligo-d(T) magnetic beads were equilibrated in 2x binding buffer (NEB, E7490L). 50 μL of RNA was then added to magnetic beads in 50 μL 2x binding buffer. Samples were incubated at 65°C on a thermocycler for 5 minutes and then the beads and RNA carefully resuspended followed by incubation at 25°C for 5 minutes, resuspension and then an additional 5-minute incubation at 25°C. The beads were captured on a magnet and the supernatant was carefully transferred to an RNase-free microcentrifuge tube. The RNA was precipitated using ammonium acetate and 100% Ethanol at −80 °C for one hour. The nascent RNA pellet was carefully dried and then resuspended in 12 μL H_2_O and stored at −80 °C for 48 hours until library preparation for sequencing. RNA quantitation and quality control were accomplished using the Bioanalyzer 2100 (Agilent Technologies). Stranded RNA-seq libraries were prepared using the KAPA RNA HyperPrep Kit with RiboErase (HMR) and KAPA Unique Dual-Indexed Adapters (Roche) according to instructions provided by the manufacturer. The barcoded paired-end libraries were sequenced by Novogene using a NovaSeq S4 generating ~70 million reads per replicate.

---

**Supplemental Figure 1.**
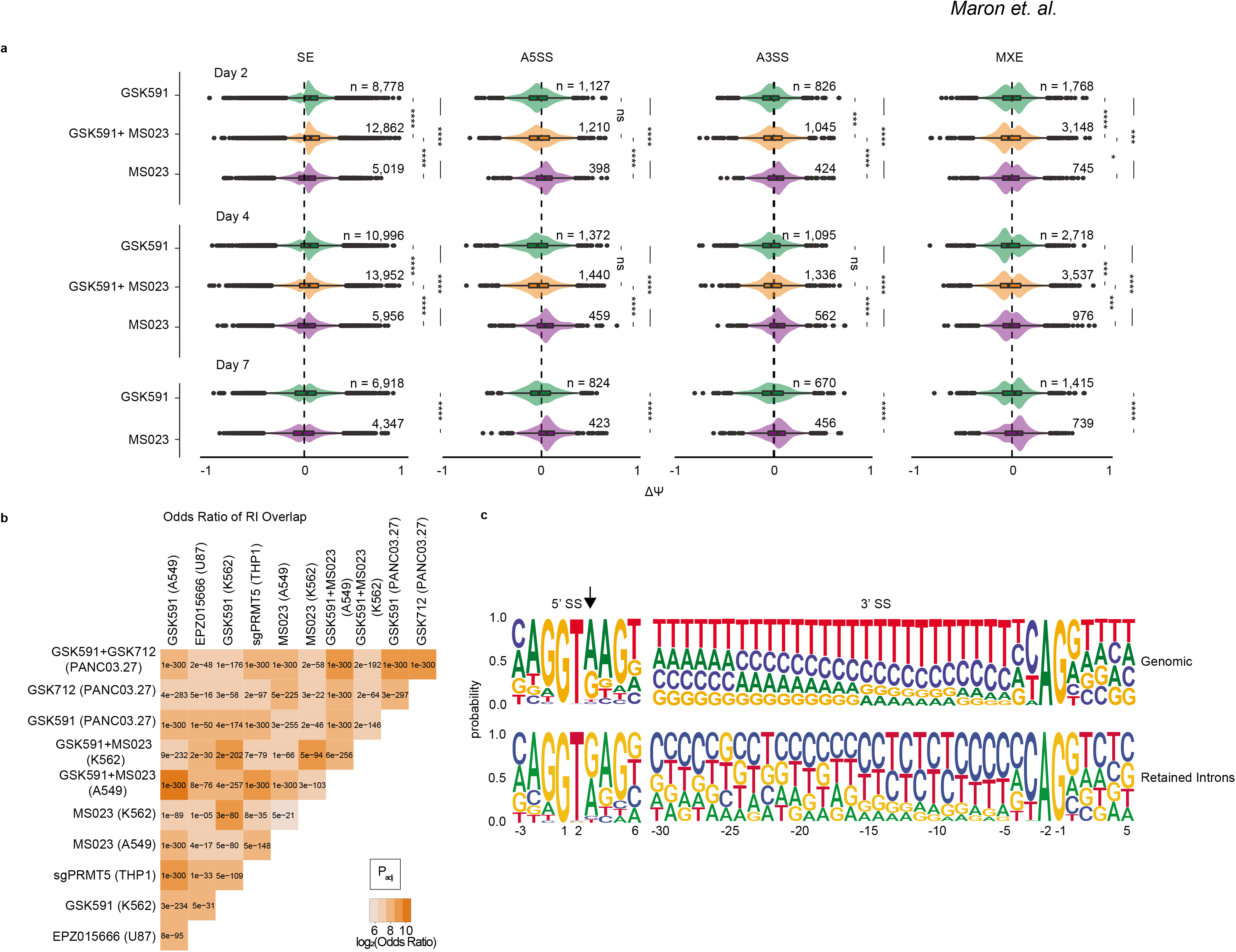
Type I PRMT or PRMT5 inhibition promotes gross changes in splicing. **a.** Comparison of the ΔΨ where ΔΨ = Ψ (DMSO) – Ψ (PRMTi) of MS023 or GSK591 treated cells at indicated time points. SE = skipped exons; A5SS = alternative 5’ splice site; A3SS = alternative 3’ splice site; MXE = mutually exclusive exons. Significance determined using two-sided Kolmogorov-Smirnov Test (* < 0.05, ** < 0.01, *** < 0.001, **** < 0.0001, ns = not significant). **b.** Matrix comparing the log_2_(odds ratio) and significance as determined by the Fisher’s Exact Test of overlapping RI from indicated experimental models. **c.** Web logo diagram of nucleotide distribution probability of global or retained introns.

**Supplemental Figure 2.**
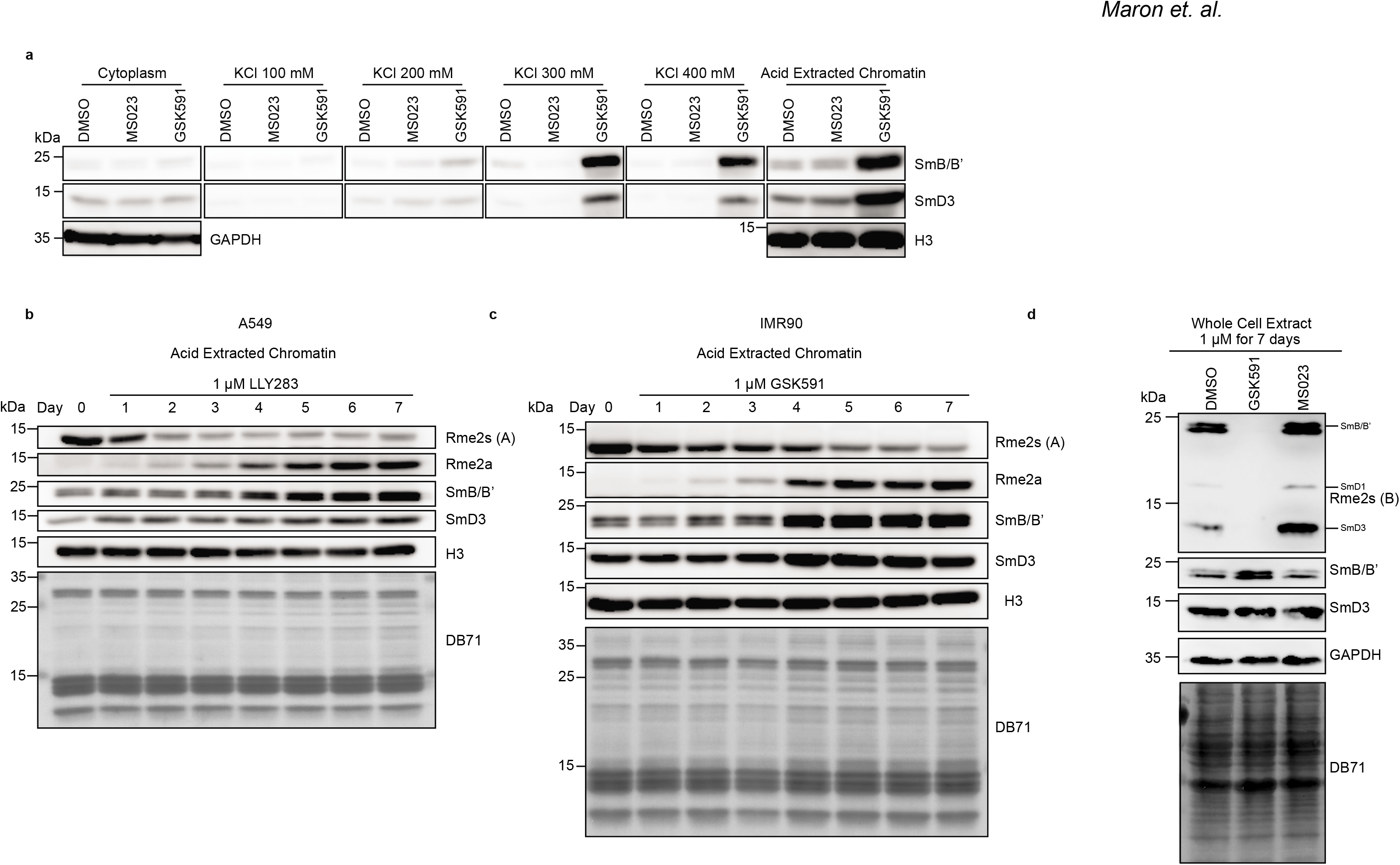
Loss of Rme2s is associated with Sm chromatin-retention. **a-d.** Western blots of hypotonic or sequential salt extraction of chromatin with indicated KCl concentrations in A549 cells treated with DMSO, GSK591, or MS023 for seven days **(a)**, acid extracted chromatin from A549 cells treated with LLY283 for the indicated duration **(b)**, acid extracted chromatin from IMR90 cells treated with GSK591 for indicated duration **(c)**, whole cell extract from A549 cells treated with PRMTi or DMSO for seven days **(d)**.

**Supplemental Figure 3.**
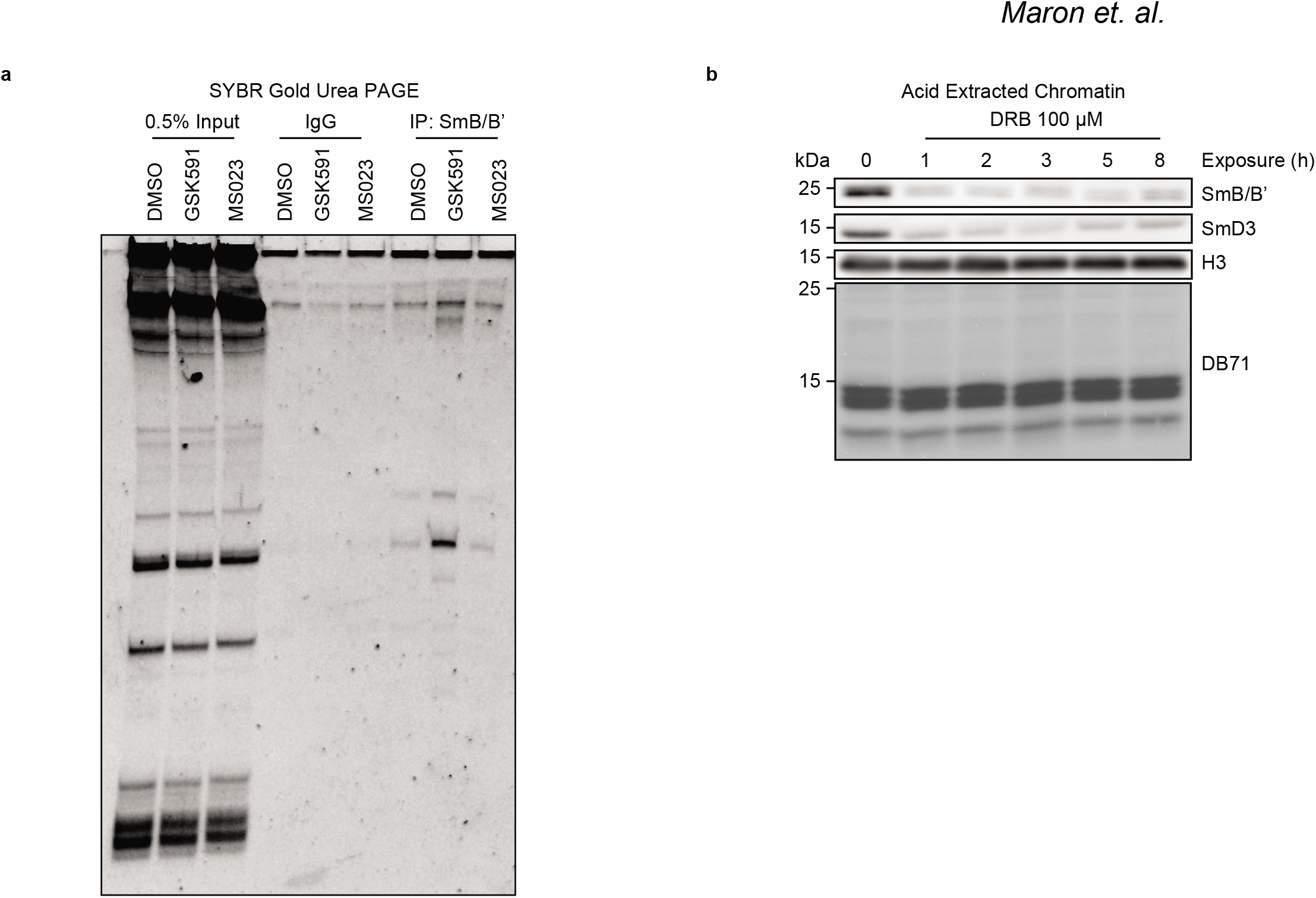
Sm proteins in PRMT5 inhibited cells are enriched for snRNA and chromatinaccumulation is transcription dependent. **a.** Urea-PAGE with SYBR gold staining of RNA immunoprecipitation with SmB/B’ in cells treated for seven-days with DMSO, GSK591, or MS023. **b.** Western blot of acid extracted chromatin from A549 cells treated with DRB for indicated time.

**Supplemental Figure 4.**
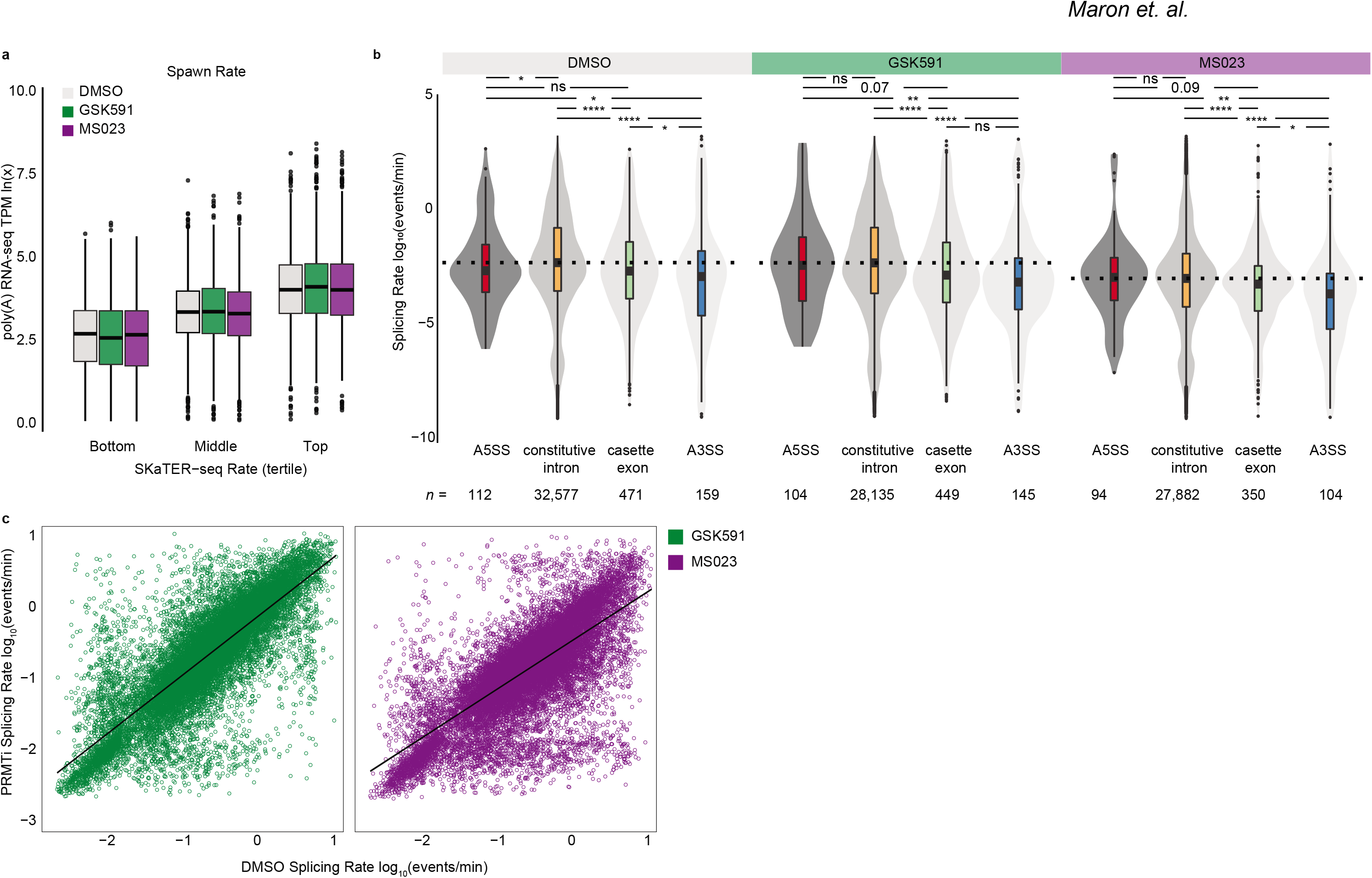
Comaprison of spawn rate to transcript abundance and splicing rate to event type. **a.** Correlation of RNA pol II initiation and pause-release (spawn rate) with poly(A)-RNA seq transcripts per million (TPM). x-axis is spawn rate in tertiles; y-axis is poly(A)-RNA seq TPM. **b.** Distribution of splicing rate versus splicing event as determined by SKaTER-seq. A5SS = alternative 5’ splice site, A3SS = alternative 3’ splice site. Significance determined using two-sided Kolmogorov-Smirnov Test (ns = not significant, * < 0.05, ** < 0.01, *** < 0.001, **** < 0.0001). **c.** Correlation of splicing rates in A549 cells treated with GSK591 or MS023 to the splicing rates in DMSO treated cells.

**Supplemental Figure 5.**
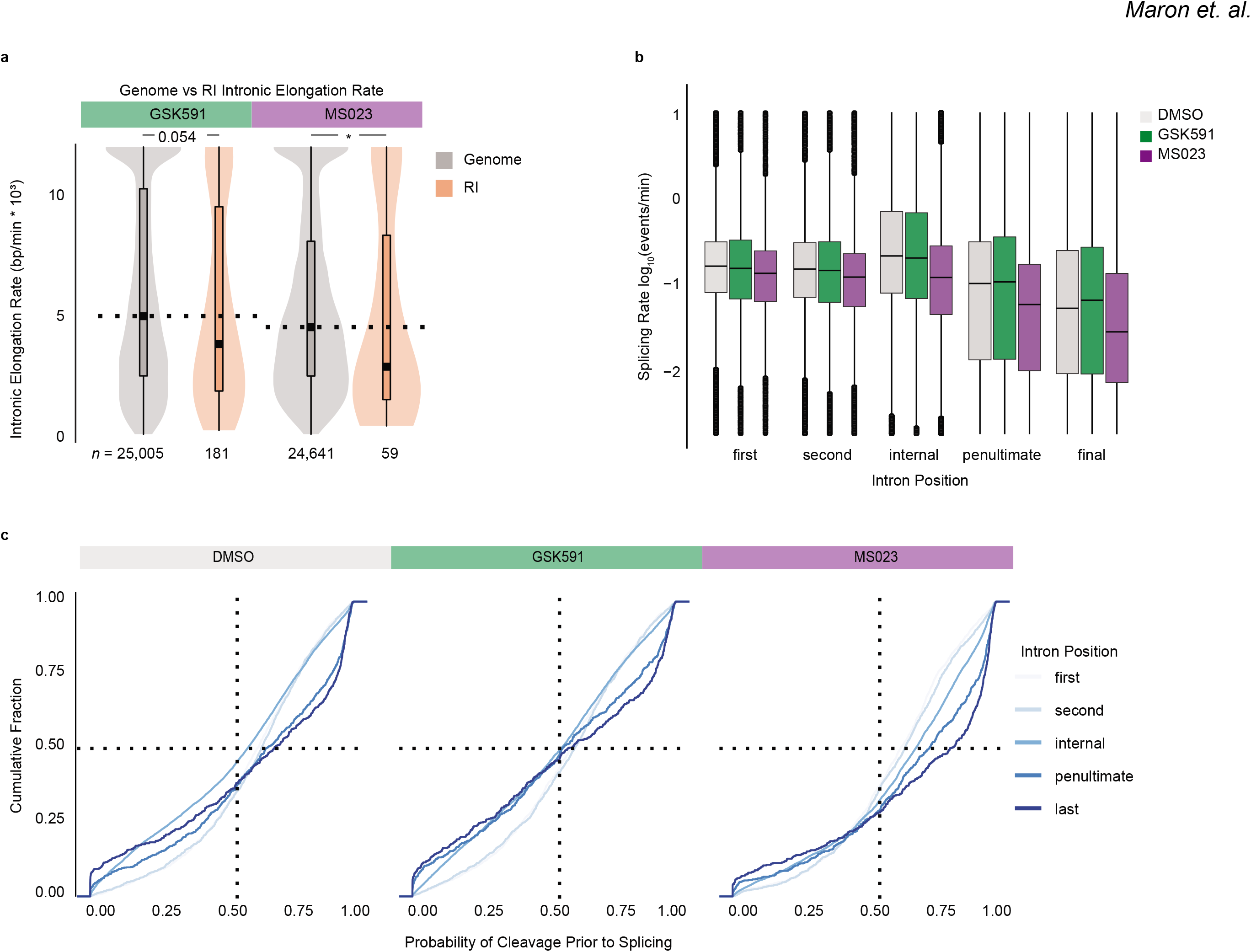
Retained introns have slower RNA pol II elongation rates and intron position correlates with splicing rate as well as the probability of cleavage prior to splicing. **a.** Comparison of RNA pol II intronic elongation rate (bp/min) in genomic versus retained introns. Significance determined using one-sided Kolmogorov-Smirnov Test (* < 0.05). **b.** Correlation of splicing rate and intron position in cells treated with DMSO, GSK591, or MS023 for two-days. **c.** Cumulative Distribution Functions of the probability of transcript cleavage prior to intron splicing compared to intron position. Color range indicates intron position where light blue is closer to TSS and dark blue closer to TES.

## Notes

### Competing Interest Statement

The authors have declared no competing interest.

